# Hierarchical Semi-Markov Smooth Models of Latent Neural States

**DOI:** 10.64898/2025.12.25.696483

**Authors:** Joshua Krause, Jacolien van Rij, Jelmer P. Borst

## Abstract

Hidden (semi-) Markov Models (HsMMs) are increasingly being used to segment neurophysiological signals into sequences of latent cognitive processes. The idea: different processes will leave distinct traces in trial-level recordings of (multivariate) neuro-physiological signals. Markov models, equipped with an *emission model* of these traces and a *latent process model* describing the progression through the different latent processes involved in a task, can then be used to infer the most likely process for any time-point and trial. However, the currently used HsMMs remain limited in two important ways. First, they cannot account for subject-level heterogeneity in the latent and emission process. Instead, a single group-level model is assumed to explain the entire data. Second, they cannot account for the potentially non-linear effects of experimental covariates on the latent and emission process. To address these problems, we present a modeling framework in which the HsMM parameters of the emission and latent process are replaced with mixed additive models, including smooth functions of experimental covariates and random effects. We derive all necessary quantities for empirical Bayes and fully Bayesian inference for all parameters and provide a Python implementation of all estimation algorithms. To demonstrate the advantages offered by this framework, we apply such a multi-level model to an existing lexical decision dataset. We show that, even in such a simple task, not all subjects rely on the same processes equally and that at least two semi-Markov states, previously believed to reflect distinct processes, might actually relate to the same cognitive process.

## 1 Introduction

Scientists have long wanted to recover the cognitive processes involved in task completion, with early attempts tracing back to Sternberg (1969) and Donders (1868). One challenge is that even simple tasks can conceivably be solved in many different ways. Successfully completing a lexical decision task for example, requiring participants to decide whether a presented character string corresponds to a word or not, implies completion of *at least* two distinct cognitive processes: initial string encoding and response initialization. Before participants can initialize the response, however, they will first have to weigh the available evidence such as the string’s orthographic plausibility and general similarity to other well-known words. However, exactly which information ultimately drives the decision, and whether this information is evaluated in a single overall decision process or in a sequence of decision-related processes, remains up for debate (e.g., Berberyan et al., 2021; Krause et al., 2024; Mahé et al., 2015; Ratcliff et al., 2004; Wagenmakers et al., 2008). This problem extends beyond lexical decision making to most cognitive tasks, with theories often disagreeing on the nature of the cognitive processes involved: their number, content, and on whether they have to be completed sequentially or can overlap at least partially (e.g., McClelland, 1979).

In this paper we present a novel modeling framework, based on hierarchical Hidden semi-Markov Models, that can be used to decode cognitive processes from time-varying neuro-physiological signals and thus help to find answers to these and related questions. The idea to decode processes from experimental data is far from new. Early attempts, which largely relied on manipulations of the experimental design and involved models of behavioral data such as reaction time (RT) measurements, date back to Sternberg (1969) and Donders (1868). Because behavioral data only provides information about the combined duration of processes required to achieve task performance, researchers recently started to rely on neuro-physiological signals to get more direct access to the processing time-course (see Anderson et al., 2016, for more detailed discussions).

The basic premise of these approaches – including the one presented here – is that different processes either rely on different neural substrates altogether or utilize them differently (for example to perform different computations), and thus leave distinct traces in neuro-physiological signals (e.g., Anderson et al., 2016; Borst & Anderson, 2015; Coquelet et al., 2022; Pieramico et al., 2025; Quinn et al., 2018; Vidaurre et al., 2016). Note, that *process* here does not necessarily refer to a single (type of) cognitive computation but also applies to *mixtures* of temporally overlapping computations associated with a distinct neuro-physiological trace (e.g., Baker et al., 2014; Pieramico et al., 2025; Vidaurre et al., 2025). Machine learning models can then be used to detect these traces and to infer the process most likely to have generated them. Recent attempts to decode processes have for example involved models of the BOLD response in different cortical regions (e.g., Anderson & Fincham, 2014; Anderson et al., 2016; Dang et al., 2017; Vidaurre et al., 2016, 2025), or multi-electrode EEG or MEG recordings (e.g., Anderson et al., 2016; Borst & Anderson, 2015; Coquelet et al., 2022; Krause et al., 2024; Lehmann et al., 1998; Vidaurre et al., 2016, 2018; Weindel et al., 2024).

Successfully decoding processes requires algorithms or methods that can reliably detect evidence of these processes in continuous time-varying signals. Conceptually, Hidden (semi) Markov Models (H(s)MMs) are ideally suited for this task. These models are characterized by two processes: a *latent process*, represented by a discrete Markov chain, and an *emission process*, producing observable output dependent on the current latent state of the Markov chain (e.g., Rabiner, 1990; Yu, 2010; Yu & Kobayashi, 2006, see also Figure 1).

**Figure 1.**
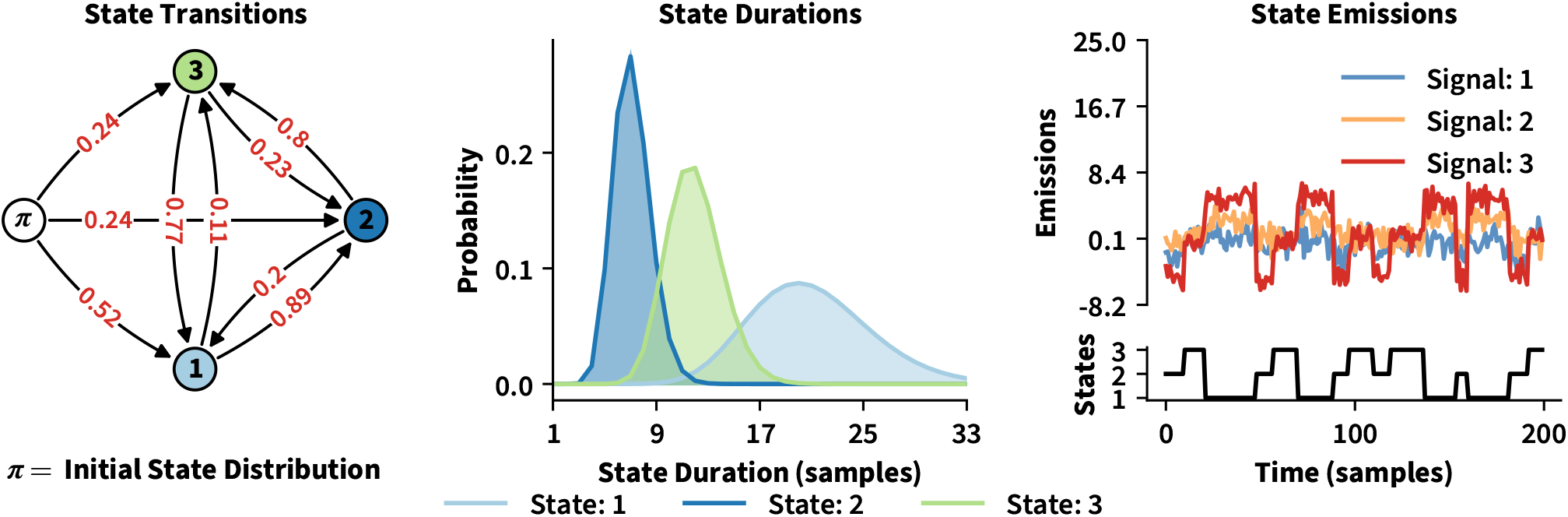
Hidden Semi-Markov Model Overview. Figure 1 visualizes the properties of the *latent* and *emission* processes characterizing a Hidden Semi Markov Model – here assumed to have three latent states. The left-most panel shows the transition graph implied by the *state transition probabilities*. Note, that self-transitions are considered impossible in many HsMMs and thus not visualized here. To account for prolonged stays in a given state, HSMMs introduce explicit *state sojourn distributions* (e.g., Yu, 2010). State-specific Gamma distributions, as visualized in the central panel, are a popular choice to model these distributions. The panel on the right shows how the latent process relates to the emission process. The panel on the bottom-right shows a possible realization of the Markov chain parameterized by the aforementioned state transition and duration probabilities. While in a given state, observable emissions are generated according to the state’s emission distribution (top-right). For this example, each state’s emission distribution was chosen as a multivariate Gaussian with distinct mean vectors and covariance matrices – in practice, differences between the emissions of different states are often more subtle.

The Markov chain can be considered a model of the potential progression through unobservable cognitive processes: just like a participant progresses through a sequence of processes to solve a certain task, the chain progresses through a sequence of distinct states according to an initial probability distribution and state transition probabilities (see left-most panel in Figure 1). As discussed in more detail in the upcoming methods section, the time spent in each state is typically modeled explicitly in Hidden semi-Markov Models – allowing these models to account for the fact that the duration of cognitive processes often varies considerably depending on the purpose they fulfill (see central panel in Figure 1).

Similarly, the emission process can be considered a formalization of the *linking hypothesis*, detailing how residence in a particular cognitive process manifests in the observable neurological signal (Anderson et al., 2016; Vidaurre et al., 2016, 2018). Conveniently, as shown in the right-most panel of Figure 1, the emission process can readily accommodate the multi-dimensional nature of most neuro-physiological data.

Another advantage offered by these models is that well-tested algorithms exist to estimate the parameters involved in the emission and latent processes from collected data. Subsequently, these parameter estimates can be used to decode the most likely realization of the Markov chain for the given data (see Yu, 2010; Zucchini et al., 2017, for an overview). Considering the conceptual parsimony and theoretical maturity of Markov models, it is thus perhaps unsurprising that they are popular among researchers wanting to recover processing stages from neuro-physiological data (e.g., Anderson et al., 2016; Borst & Anderson, 2015; Coquelet et al., 2022; Dang et al., 2017; Krause et al., 2024; Pieramico et al., 2025; Vidaurre et al., 2016, 2018, 2025; Weindel et al., 2024).

However, in contrast to the models presented in this paper, the specific (Semi) Markov models currently used to recover processing stages remain limited in two important ways. First, these models typically do not account directly for subject-level heterogeneity in the latent and emission process. Second, they cannot readily account for potentially non-linear effects of experimental covariates on the latent and emission processes.

Specifically, while data is commonly collected from different subjects, all of whom complete multiple experimental trials, it is common to estimate only a single set of parameters applicable to the entire group. While such a group-level model can still be used to decode the separate (order and duration of) processes completed on individual trials, doing so assumes that the time-series from individual trials reflect different realizations of the same Markov chain and emission process. This assumption is difficult to justify, since different subjects can generally be expected to vary systematically in the time it takes them to complete individual cognitive processes.

Similarly, some participants might simply not complete a specific process at all or might gradually stop to do so because they become more familiar with the task or adapt their strategy over time (e.g., Baayen et al., 2017). Even if data were to be collected from a single expert participant, any experimental covariate designed to influence task difficulty would almost certainly influence the properties of the latent process as well – and might do so non-linearly. Indeed, in our recent work we observed strong non-linear effects of word-likeness on the duration of virtually all processes involved in lexical decision making (Krause et al., 2024). There we also discussed the possibility of word-likeness influencing the neural pattern indicating residence in a particular cognitive process, and that it is plausible that different subjects might deviate differently from this pattern as well.

In summary, subject-level heterogeneity and non-linear covariate effects both introduce dependencies into the emission and latent process, which current models cannot readily account for. To address these dependencies we propose to replace constant group-level parameters of HsMMs with additive mixed (or multi-level) models of the parameters (or a known function of them; cf. Michelot, 2025; Wood et al., 2016). The multi-level models of individual parameters can include any combination of smooth functions of experimental covariates and i.i.d random effects to capture subject-level heterogeneity.

We start by formally introducing the specific semi-Markov model used most commonly by researchers wanting to detect cognitive processes in the upcoming section 2.1. We then outline how this model can be expanded to include non-linear and random effects (section 2.2) and derive the quantities necessary to estimate the resulting models, such as the log-likelihood function and corresponding gradient (sections 2.2.1 - 2.2.3). We also provide an implementation of the discussed algorithms in the open-source Python toolbox `mssm` (version ≥1.2.0), available on GitHub and distributed via the Python Packaging Index^1^.

In section 3 we demonstrate the usefulness of the framework presented here: we present a re-analysis of the lexical decision dataset collected by Krause et al. (2024), showing that only a multi-level model accounting for subject-level heterogeneity and non-linear covariate effects could have generated the data. While the improved model generally makes similar predictions about group-level effects of word-likeness and word-type, it also reveals that not all subjects seem to rely on the same processes equally and that at least two semi-Markov states, previously believed to reflect distinct processes, might actually relate to the same cognitive process.

Finally, in section 4 we discuss the implications of the framework presented here for the task of *model selection*: identifying the model most likely to have generated the data from a set of candidates. The space of possible models grows substantially once random effects and smooth functions of covariates are taken into account, which significantly complicates this task. We discuss different options to address this problem and their respective limitations. Finally, we discuss possible extensions and applications of the framework presented here.

## 2 Methods

Recent attempts to recover processing stages mostly rely on some kind of Hidden (semi-) Markov model. The particular semi-Markov model most commonly used is the “Explicit Duration Hidden Markov Model” (EDHMM), which is a specific type of HsMM (see Levinson, 1986; Rabiner, 1990; Yu, 2010). In the next section we conduct a formal review of this type of model. To simplify notation for this review, we assume that only a single time-series is observed.

In section 2.1.3 we outline how the EDHMM can be generalized to account for multiple time-series (see also Rabiner, 1990).

### 2.1 Review: Explicit Duration Hidden Markov Models

Like other HMMs (and HsMMs), the EDHMM is characterized by a *latent process*, taking on the form of a discrete Markov chain of latent states 𝒮 ={1, …, *M*}, and an *emission process*, producing observable output *o*_*t*_ (e.g., EEG amplitudes or BOLD responses) dependent on the current state *S* _*t*_ of the Markov chain (e.g., Rabiner, 1990; Yu, 2010; Yu & Kobayashi, 2006). The main difference between HsMMs, including the EDHMM, and conventional HMMs, is that HsMMs also include an explicit model of the time spent in each state as part of the latent process model (e.g., Rabiner, 1990; Yu, 2010). In the next two sub-sections we provide a formal review of the properties of the latent and emission process of an EDHMM.

#### 2.1.1 The Latent Process of an EDHMM

The latent process of an EDHMM is a Markov chain with the extension that time spent in each state is modeled via explicit distributions (e.g., Rabiner, 1990; Yu, 2010). We first consider the properties of the Markov chain, which is defined by a probability mass function over the initial states and state transition probabilities (Yu, 2010). It is assumed that at *t* = 1, the Markov chain is in state *j* with probability 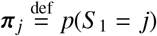. *t* by convention denotes the current discrete time-point. In the context of this paper, it thus makes sense to use *t* to refer to a particular sample of neuro-physiological data. For example, *t* = 1 will typically reflect the sample coinciding with the onset of a trial or stimulus. Similarly, the last observable output *o*_*T*_ is generated by state *S* _*T*_ at time-point *t* = *T*, which in the context of this paper typically reflects the last sample collected during a trial (for example when a participant gives a response).

To simplify the computation of the likelihood, it is conventionally assumed that a new state *begins* at *t* = 1 – in contrast to continuing from *t* = 0, *t* = −1, etc – and that the final state ends at *t* = *T*, after omitting the final emission (e.g., Yu, 2010). This assumption is not always easy to justify when latent states are assumed to correspond to cognitive processes. For example, in some experiments participants might come to anticipate the onset of the next stimulus and it would be unreasonable to assume that the onset of the first cognitive process coincides with stimulus onset. We return to these “censoring” issues, and how they can be addressed in section 2.2.1 (see also Yu, 2010, for an overview of different censoring issues).

The chain then repeatedly transitions from one state to another state according to the *state transition probabilities* (see left-most panel of Figure 1). We denote the transition from state *i* to state *j* with 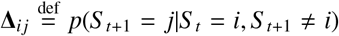 – all collected in the *M M* matrix Δ. For conventional HMMs, self-transitions are the mechanism to account for prolonged stays in a given state. When working with HsMMs, the time spent in each state (i.e., the “state sojourn time “; Yu, 2010) is modeled explicitly instead. As such, self-transitions are considered impossible by convention for the EDHMM, so that Δ_*ii*_ = 0 ∀*i* ∈*S* (cf. Yu, 2010).

The state sojourn time is treated as a discrete random variable *d* with state-dependent probability mass function *p*_*j*_(*d*) for *j* ∈𝒮 and *d* ∈{1, …, *D*} (e.g., Levinson, 1986; Yu, 2010; Yu & Kobayashi, 2006). Alternatively, a (normalized) probability density function can be used for the model of the sojourn time instead (e.g., Levinson, 1986; Yu, 2010). That is, *d* | *S* = *j* ∼ ℱ^*d*^(ν _*j*_, …, *γ*_*j*_), where ℱ^*d*^ is a state-specific duration or state sojourn time distribution parameterized by ν _*j*_, …, *γ*_*j*_. This drastically lowers the number of parameters required in the model of the durations expected for each state. While different distributions may require a different number of parameters, a mean ν _*j*_ parameter and optional scale parameter *γ*_*j*_ are sufficient when relying on a (normalized) member of the exponential family for ℱ^*d*^ (see Yu, 2010, and the central panel of Figure 1).

#### 2.1.2 The Emission Process of an EDHMM

The output *o*_*t*_ observed at time-point *t* is assumed to depend on the current state of the chain (e.g., Yu, 2010). Conventionally, the EDHMM assumes univariate emissions, believed to be independent and identically distributed (i.i.d) given knowledge about the current state of the system (this “conditional independence” assumption is common when working with other H(s)MMs as well, see; Levinson, 1986; Rabiner, 1990; Yu, 2010; Yu & Kobayashi, 2006). Hence, *o*_*t*_ | *S* _*t*_ = *j* ∼ ℱ^*o*^(*µ*_*j*_, …, *ϕ*_*j*_), where ℱ^*o*^ denotes a state-specific *emission distribution* or *emission model*. Common choices for ℱ^*o*^ are again members of the exponential family, such as Gaussian or Gamma distributions (e.g., Yu, 2010), which only require a mean *µ*_*j*_ and optional scale parameter *ϕ*_*j*_.

When using EDHMMs to segment neuro-physiological signals into cognitive processes, it is typically necessary to account for multivariate output (e.g., a *K* vector of EEG amplitudes recorded from multiple electrodes). Multiple strategies exist to ensure that the EDHMM generalizes to multi-variate output.

A popular option is to assume that the multivariate output vectors **o**_*t*_ are generated by state-specific multivariate normal distributions with mean vectors *µ*_*j*_ and covariance matrices Σ _*j*_ – which also permits for multivariate auto-regressive emission models (e.g., Vidaurre et al., 2016). Such emissions models have been applied to trial-level EEG and MEG data directly (e.g., Vidaurre et al., 2016, 2018). More recently however, researchers have started to first transform the raw data, creating “virtual channels”, sometimes also called “delayed embeddings”, that carry information about temporal and spatial relations between and within raw data channels (i.e., electrodes) (see Borst & Anderson, 2015; Vidaurre et al., 2018, 2025). Subsequently, a Principal Component Analysis (PCA) is applied to the transformed data and a multi-variate Gaussian emission model is assumed for the *K* top components (e.g., Vidaurre et al., 2018, 2025).

Alternatively, it is common to treat the emissions of the *K* signals as conditionally independent, given knowledge about the current state (e.g., Anderson et al., 2016; Borst & Anderson, 2015; Weindel et al., 2024). This is essentially a generalization of the univariate emission model outlined earlier. Specifically, one assumes that 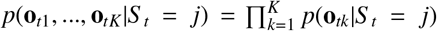. *p*(**o**_*tk*_|*S* _*t*_ = *j*) denotes the density implied by the conditional univariate distribution **o**_*tk*_ *S* _*t*_ = *j* ∼ ℱ^*o*^(*µ*_*jk*_, …, *ϕ*_*jk*_), where **o**_*tk*_ is the emission generated by signal *k* at time-point *t* (i.e., element *k* of emission vector **o**_*t*_, see also; Zucchini et al., 2017). Anderson et al. (2016) rely on this assumption for example, to derive their “multivariate pattern” emission model of EEG data. This model of the emissions assumes that the onset of a new process is assumed to result in the addition of a weighted half-sine of fixed duration to the EEG signal at each electrode, which can otherwise be described as a multivariate oscillatory noise process with zero expectation. To estimate an EDHMM, the authors compute a PCA of the raw EEG data and subsequently treat the *K* top components as independent (see Appendix A for more details, but also Anderson et al., 2016).

To avoid notational clutter we write **o**_*t*_ from now on, for both univariate and multivariate emissions and independent of whether a multivariate Gaussian or multiple independent densities are used as the model of multivariate emissions.

#### 2.1.3 Multiple Time-Series, Subject-level Differences, and Covariate Effects

The EDHMM outlined above can account for data collected from multiple time-series and for possible trial-level variability (also between conditions or subjects) in the onset of cognitive processes – if we assume that different time-series reflect independent realizations of the *same* Markov chain and emission process (e.g., Zucchini et al., 2017). This assumption is attractive because only a single set of *group-level* parameters applicable to all collected time-series, subjects, etc. has to be estimated. Additionally, the independence assumption drastically simplifies computation of the model’s likelihood (e.g., Yu, 2010; Zucchini et al., 2017).

While it is common to assume independence of different time-series given group-level parameter estimates only, this is not justified when dealing with the problem of recovering processing stages from neuro-physiological data due to the reasons outlined in the introduction: All of the aforementioned emission models assume a distinct neural pattern of some form, indicating the onset of or residence in a particular cognitive process, and different subjects might deviate from this pattern differently. Additionally, experimental covariates (e.g., any continuous or trial-specific manipulation of task difficulty) are likely to influence this pattern as well – and they might do so non-linearly.

Similarly, it is all but guaranteed that there will be systematic differences between subjects in the timing and the order of the underlying cognitive processes, i.e., the latent process. Finally, experimental covariates are just as likely to affect the properties of the latent process, as they are to affect the pattern in the observable data.

Previous work typically addresses the possibility of subject-level differences and non-linear covariate effects with a two-step or dual estimation process (e.g., Krause et al., 2024; Vidaurre et al., 2025). In a first step, group-level parameters of the EDHMM are estimated. These parameters are used to decode the most probable state sequence for every subject. In a second estimation step, subject-specific EDHMM parameters and functions of covariates are estimated given the recovered state sequences, which are kept fixed. While effective, such procedures are limited by the fact that there is no guarantee that similar state sequences would be produced by a model in which the EDHMM parameters are allowed to vary between subjects and as a function of experimental covariates. As such, subject-level and function estimates obtained from group-level state sequences remain biased.

Notably, if the true number of states *M* (i.e., processes) is assumed to be known in advance, this bias might not be of practical relevance. This will depend on factors such as the degree of subject-level variability and the amount of data available to estimate the group-level models. In contrast, the bias might be far more substantial when *M* is to be estimated alongside the Markov model parameters, for example via leave-one-out cross-validation (LOOCV; e.g., Krause et al., 2024). In that case, a model including only group-level parameters might already result in a different estimate for *M* than a model in which parameters are allowed to vary. Ultimately, two-step estimates from this model could thus be vastly different from the subject-level and covariate effects produced by a model with varying parameters.

As such, it would generally be desirable to be able to estimate a model, not unlike a multi-level EDHMM, that permits non-linear covariate effects on model parameters and for these parameters to vary between subjects. Crucially, assuming independence of different time-series given model parameters would be far less problematic for such a model, since the parameters (and hence the latent and emission processes) themselves can vary between subjects and as a function of experimental covariates (e.g., Michelot, 2025). The next section outlines how such a model, which we will refer to as a *multi-level semi-Markov smooth model*, can be formalized.

### 2.2 Multi-level Semi-Markov Smooth Models

As mentioned briefly at the end of the previous section, we propose to borrow the latent and emission processes of the EDHMM but to model their definining parameters, or a known function of them, as an additive combination of smooth functions of covariates and i.i.d random effects to capture subject-level heterogeneity (cf. Wood et al., 2016). Technically, the class of multi-level semi-Markov smooth models (SMSMs) generalizes beyond the emission and latent processes assumed by an EDHMM to those assumed by any of the other common HsMMs discussed by Yu (2010). Even more generally, the SMSM framework is applicable to other latent variable models, potentially including additional latent variables related to the emission process (e.g., Taghia et al., 2018), requiring only that the gradient of the log-likelihood function can be computed. Here we consider an EDHMM-like SMSM as an example, but much of the theory will apply to SMSM extensions of other models. Additionally, the specific SMSM presented here can be considered a semi-Markov extension of the models discussed by Michelot (2025), who focused largely on conventional HMMs without explicit models of the state sojourn times^2^.

To formalize an EDHMM-like SMSM, we first consider the parameters required by emission and state sojourn time distributions. Let *µ*_*j*_ denote any parameter involved in the model of the emission process of state *j* that is required by emission distributions ℱ^*o*^. For example, *µ*_*j*_ could be a mean of a (multivariate) normal distribution, the scale parameter of a Gamma distribution, or a covariance parameter in the precision matrix of a multivariate normal distribution (i.e., an off-diagonal element in Σ^−1^). In addition, let ν _*j*_ denote any parameter required by the duration distribution ℱ^*d*^ of state *j*. For example, ν _*j*_ could be the mean of a Gamma or Poisson distribution, or the scale parameter of an inverse Gaussian distribution.

These and the remaining parameters, the state transition probabilities in Δ and initial state probabilities in *π*, are considered constant in the conventional EDHMM. In contrast, in a SMSM they are each parameterized by a *linear predictor η* (e.g., Michelot, 2025; Wood et al., 2016). Here we are interested in linear predictors of the form outlined in Equation (1).

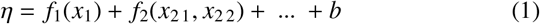

Here, *b* is a random coefficient so that 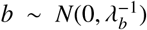 and *f*_1_ and *f*_2_ are smooth functions of covariates *x*_1_ and *x*_21_, *x*_22_ respectively. Functions like *f* are represented as weighted sums of δ known *basis functions b* so that (e.g., Krause et al., 2025; Wood, 2017; Wood et al., 2016)

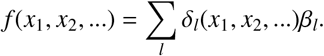

To prevent overfitting, functions like *f*_1_ and *f*_2_ are regularized by placing an (improper) *smoothness* prior on the vector of coefficients *β*_*k*_ associated with smooth function *f*_*k*_, so that 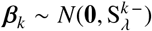 (e.g., Kimeldorf & Wahba, 1970; Wood, 2017; Wood et al., 2016). Here, 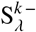 denotes the pseudo-inverse of the semi-definite precision matrix 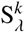, which has a known structure up to one or more scaling variables *λ* ∈*λ*_*k*_ (see Wood, 2017, for a more detailed discussion). These *λ* parameters are essentially regularization hyper-parameters with their magnitude determining how complex functions like *f* can be. Thus, they will typically have to be estimated like *β*_*k*_ – we discuss this in an upcoming section. Note also, that the prior is improper to allow treating sufficiently simple function estimates (i.e., lower-order polynomials) as fixed (see Wood, 2017).

Representing functions like *f*_1_ and *f*_2_ as sums of weighted basis functions reveals that Equation (1) matches an *additive mixed model* of the form shown in Equation (2) (see also Michelot, 2025).

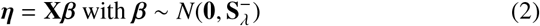

Here, **X** corresponds to the model matrix of *η* and all coefficients (e.g., *β*_*k*_, *b*, …) involved in the model of *η* have been collected in *β*. As discussed, *β* is subjected to an improper Gaussian prior, with 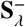 denoting the pseudo inverse of semi-2017; Wood et al., 2016). 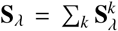 is obtained by sum-definite precision matrix **S**_*λ*_ (e.g., Krause et al., 2025; Wood, ming over embedded^3^ versions 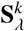 of all precision matrices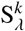 involved in the model of *η*, including those associated with standard i.i.d random coefficients (e.g., like *b* in Equation 1) for which 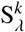 is simply a scaled identity (Wood, 2017).

Based on the definition of linear predictors in Equation (2), we define models of parameters *µ*_*j*_ and ν _*j*_ as shown in Equation (3).

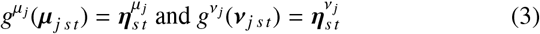

Here, *g* are known, optionally state-specific, smooth *link* functions with known inverse so that, for example, 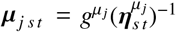 (e.g., Faraway, 2016; Wood, 2017). Subscripts indicate the value of a parameter (e.g., *µ*_*j*_) or predictor (e.g., 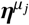) for time-series (i.e., trial) *s* and at time-point *t*. Specifically, 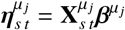, where 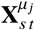 indexes the specific row in the subset of model matrix 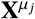 associated with parameter *µ* and state *j* for time-series *s* and time-point *t*. Finally, 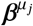 denotes the set of coefficients involved in the model of 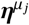.

We now turn to the state transition probabilities Δ_*ij*_. As mentioned, these are also expressed in terms of a linear predictor in a SMSM. To achieve this, first note that the offdiagonal probabilities in row *i* of matrix Δ define the mass function for a Multinomial distribution of the transitions away from state *i* to state *j* ∈𝒮*/i* (e.g., Michelot, 2025). Thus, we can write

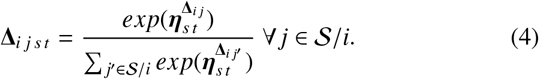

Note, that for every state *i* only *M* −2 linear predictors 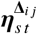 are identifiable, requiring the additional constraint that at least one of those is set to zero. The choice is essentially arbitrary and can also differ for different states *i* (see also Michelot, 2025). A similar parameterization is chosen for *π*. Specifically, the elements in *π* again form the mass function for a Multinomial distribution over the initial state. Hence, we again write:

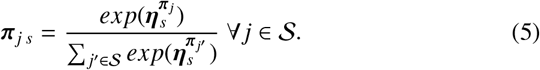

Again, only *M* − 1 linear predictors 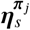 are identifiable, so we enforce the constraint that 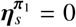. Also, note that the *t* sub-scripts have been omitted in this equation, since the initial state distribution cannot change over time.

With models specified for the different parameters as shown above, it is now possible to account for subject-level heterogeneity in the emission and latent processes as well as for non-linear covariate effects. Note, that the models of the parameters outlined above have been kept extremely general – considerable simplifications will often be possible (and even necessary, if the model is to be estimated in a reasonable amount of time). For example, many experiments will only contain covariates varying between time-series but not within (i.e., with time). In that case, the *t* sub-scripts can be omitted in either (or both) of Equations (3) and (4). Similarly, prior knowledge might be available regarding the structure and elements of Δ and *π*. In that case, models could be specified only for the parameters involved in the emission and state duration distributions (e.g., *µ*_*j*_ and ν _*j*_). The next section outlines how the log-likelihood of the specific EDHMM-like SMSM considered here can be computed efficiently.

#### 2.2.1 The Log-likelihood of the EDHMM-like Semi-Markov Smooth Model

To facilitate the likelihood definition, let *β* denote the total set of coefficients, required by the models of all parameters. Next, we show how the log-likelihood ℒ_*s*_(*β*) = *log*(*p*(**o**_*s* [1:*T*]_|*β*)) of *β* can be computed given the emission sequence 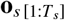 corresponding to an individual time-series *s*. We use square brackets in sub-scripts to index a sequence of elements for a given dimension (i.e., [1 : *T*_*s*_] indexes time-points 1, …, *T*_*s*_). Additionally, *T*_*s*_ has received a subscript to emphasize that the number of time-points collected can differ between series. Given a way to compute the log-likelihood for individual series, the total log-likelihood is then simply ℒ = ∑_*s*_ ℒ_*s*_(*β*), due to the assumption that different time-series are mutually independent given *β*.

As for other Markov models, ℒ_*s*_ is best computed recursively for the model considered here, utilizing the so-called *forward variables* denoted by *α* (e.g., Rabiner, 1990; Yu, 2010). The forward variables presented here are scaled versions of the ones defined previously by Yu and Kobayashi (2006). The scaling, previously introduced by Lystig and Hughes (2002) in the context of conventional HMMs, drastically facilitates computation of the gradient of the log-likelihood later on. For *t* = 1 the variables are initialized as shown in Equation (6).

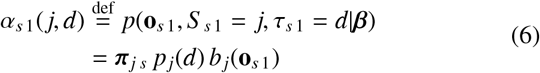

Here, *τ*_*t*_ is an auxiliary variable denoting the remaining or residual duration in the current state at time-point *t* and *π*_*j s*_ is as defined in Equation (5). Additionally, *p*_*j*_(*d*) and *b*_*j*_(**o**_*s t*_) are again used as shorthands for 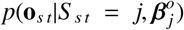 and 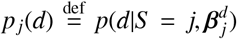 respectively, where 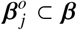 and 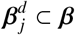 denote the sets of coefficients required to parameterize the emission and state sojourn time distributions ℱ^*o*^ and ℱ^*d*^ of state *j* ^4^. Note that conditioning on 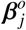 and 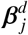 implicitly defines *t* and *s* subscripts for the densities/probabilities *p*_*j*_(*d*) and *b*_*j*_(**o**_*s t*_), since the values of parameters required by ℱ^*o*^ and ℱ^*d*^ can change for different series and time-points as shown in Equation (3).

For *T*_*s*_ ≥*t* ≥2, the forward variables can then be computed recursively as shown in Equation (7).

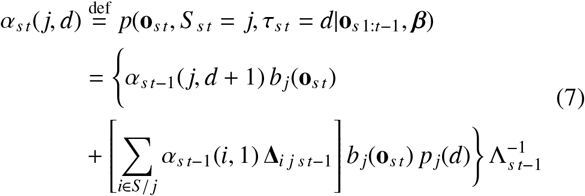

Here Δ_*i j s t*−1_ is defined as in Equation (4) and

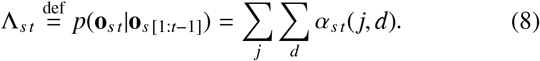

Intuitively, the forward variable defined in Equation (7) thus computes the desired probability *p*(**o**_*s t*_, *S* _*s t*_ = *j, τ*_*s t*_ = *d*|**o**_*s* [1:*t*−1]_, *β*) based on the probability of the same state *j* continuing (i.e., having duration *τ*_*s t*−1_ = *d* + 1 at the previous time-point *t* −1) and the probability of any state *i* ∈*S/ j* ending on the previous time-point (i.e., having remaining duration *τ*_*s t*−1_ = 1) with the chain then transitioning from state *i* into *j* for duration *d* (Yu & Kobayashi, 2006).

Note, that some forward variables can be set to zero and thus do not have to be computed, in case the possibility of right-censoring can be discounted (e.g., Yu, 2010). Specifically, if there is prior knowledge suggesting that the last state ends at the last time-point *t* = *T*, we can set *α*_*s t*_(*j, d*) = 0 ∀*t* + *d > T*_*s*_ + 1. Similarly, if there is prior knowledge suggesting that the last state is *j*, we can set 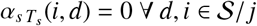.

Accounting for left-censoring (e.g., Yu, 2010) is more complicated. Specifically, use of *p*_*j*_(*d*) in both Equations (6) and (7) implies the assumption that the first state begins at the first time-point *t* = 1 (i.e., that there is no left-censoring). If this is not the case, then the mass associated with duration *d* at *t* = 1 could be very different than at other time-points, and use of *p*_*j*_(*d*) in both Equations will no longer be justified. To this end we propose to generally^5^ replace *p*_*j*_(*d*) with *p*_*π j*_(*d*) in Equation (6). *p*_*π j*_(*d*), like *p*_*j*_(*d*), is parameterized as outlined in Equation (3), but comes with a separate set of coefficients 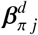. If there is prior evidence suggesting that the possibility of left-censoring can be discounted, we can set 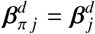. Otherwise these extra coefficients will have to be estimated (see also Yu, 2010).

Finally, note that the definition of the forward variables given in Equations (6) and (7) also permits emission models for which the density *p*(**o**_*s t*_ | *S* _*s t*_ = *j*) cannot easily be computed, but *p*(**o**_*s t*_ | *S* _*s t*_ = *j, τ*_*s t*_ = *d*) can. As discussed in more detail in Appendix A, this is necessary for example for the emission model of EEG proposed by Anderson et al. (2016). With the forward variables defined, the log-likelihood of the EDHMM-like latent smooth model for series *s* can then be computed based on the Λ_*s t*_ variables defined in Equation (8). Specifically, relying on the fact that any likelihood can always be factorized into a product of conditional probabilities (e.g., Lystig & Hughes, 2002),

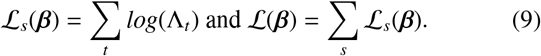

Note, that while the computational complexity of the forward variables is considerable, Equations (6) and (7) can be evaluated in parallel for different series *s*, which can drastically speed up the computation of (*β*). The forward variables are also useful for a broad range of other tasks. For example, they are required to compute the most likely state sequence, given estimates of *β*, for every series via the Viterbi algorithm (e.g., Rabiner, 1990). Similarly, they are required for sampling state sequences given data and estimates of *β* (e.g., Dewar et al., 2012). The next section considers how to obtain such estimates of *β* for a SMSM.

#### 2.2.2 Estimating Semi-Markov Smooth Models

The main challenge complicating estimation of SMSMs is that the (inverted) variance/regularization parameters *λ*, involved in the priors placed on random effects and smooth functions *f* in any of the models defined in Equations (3-5), will have to be estimated as well. Here we consider two possible solutions to this problem: an empirical Bayes approach and fully Bayesian inference.

The *empirical approach* is to (approximately) optimize the Bayesian marginal likelihood *p*(**O** | *λ*) (e.g., Krause et al., 2025; Wahba, 1985; Wood, 2017; Wood et al., 2016). Here we use **O** to denote the concatenated observation sequences from all time-series and recorded signals (i.e., all the data) and *λ* again denotes the set of all variance/regularization parameters required by the model. Let *p*(*β*|*λ*) denote the density implied by the (improper) prior Gaussian prior 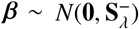 placed on the total coefficient vector *β*. Note, that 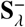 now denotes the block-diagonal matrix obtained by stacking the 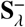 matrices defined for each linear predictor (see Equation 2; and also Krause et al., 2025; Wood et al., 2016, for a more detailed description). Then, the Bayesian marginal likelihood can be computed (approximately) as shown in Equation (10).

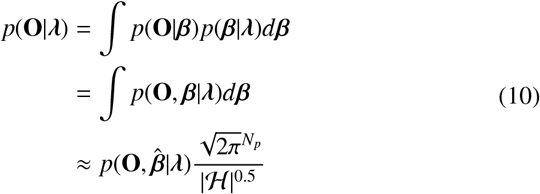

Here, *p*(**O** *β*) = (*β*), *N*_*p*_ is the number of elements in *β*, and the approximate result of the third line follows from a Laplace approximation to the conditional posterior *β* **O**, *λ* (see Wood, 2017; Wood et al., 2016, for more detailed discussions). 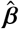 is the maximizer of the joint log-likelihood *log*(*p*(**O**, *β λ*)). Similarly, ℋ is the negative Hessian of the joint log-likelihood, evaluated at said maximizer. Considering that the density of *p*(*β λ*) is a zero-mean multivariate Gaussian, it can readily be shown that 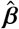 is obtained by maximizing the *penalized* log-likelihood 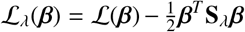 while H = **H** + **S**, where 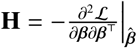 is the negative Hessian of the log-likelihood evaluated at 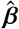 (e.g., Krause et al., 2025; Wood et al., 2016).

Maximizing *log*(*p*(**O**|*λ*)), or the Laplace approximate version V(*λ*) = *log*(*p*_*L*_(**O**|*λ*)) given in the third line of Equation (10), thus automatically provides point-estimates 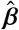 along-side point-estimates 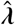 (e.g., Wood, 2017). Additionally, the Laplace approximation provides a normal approximation to the conditional posterior 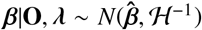 (Wood, 2017; Wood et al., 2016). Combined with the asymptotic approximation 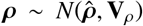, where *ρ* = *log*(*λ*) and 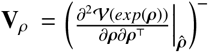, this enables approximate inference about *β* and *λ* (Krause et al., 2025; Wood, 2017; Wood et al., 2016).

Different approaches have been proposed to maximize 𝒱 (*λ*) numerically. Wood et al. (2016) outline how exact gradients and the hessian of 𝒱 (*λ*) can be computed. Both, ℒ_*λ*_(*β*) and 𝒱 (*λ*) can then be maximized via Newton’s method. However, this approach generally requires up to fourth order derivatives of the log-likelihood ℒ. In contrast, the Extended Fellner Schall (EFS) method described by Wood and Fasiolo (2017) only requires the gradient of the log-likelihood and **H** (i.e., up to second-order derivatives) to optimize ℒ_*λ*_(*β*) via Newton’s method and to approximately maximize 𝒱 (*λ*).

Michelot (2025) show that automatic differentiation of the computer code of the log-likelihood can be used to obtain the gradient and **H**, again enabling optimization of _*λ*_(*β*) and computation of (*λ*). The latter can then again be maximized based on the EFS method or via quasi-Newton methods relying on numerical approximations to the derivatives of (*λ*) (see Michelot, 2025; Nocedal & Wright, 2006). Any computations involving the negative Hessian **H** will however become expensive for complex models involving many random effects. To address this problem, Krause et al. (2025) recently showed that **H** in the EFS update can also be replaced with a limited-memory symmetric rank one approximation, computed from accumulated gradients of the log-likelihood and modified to remain semi-definite (see also Nocedal & Wright, 2006). The resulting limited-memory quasi-EFS (L-qEFS) update thus avoids explicitly forming **H**, and only requires the gradient of to approximately optimize _*λ*_(*β*), via quasi-Newton methods, and (*λ*) via the modified EFS update.

While the gradient of the log-likelihood ℒ (*β*) of a SMSM could be derived via automatic differentiation (e.g., Michelot, 2025), an analytic solution can be computed recursively for the specific EDHMM-like model considered here, just like the forward variables defined in Equations (6) and (7). The analytic solution was previously derived by Lystig and Hughes (2002) for conventional HMMs – here we generalize their approach to latent smooth models. First, Lystig and Hughes (2002) point out that

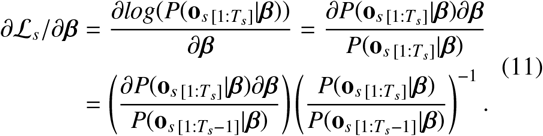

Based on the definitions given in Equation (8),

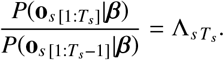

Crucially, Lystig and Hughes (2002) then point out that the remaining quantity in Equation (11), 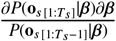, can be computed based on the partial derivatives of the forward variables with respect to *β*. To achieve this for the specific definitions of the forward variables shown in Equations (6) and (6), we first initialize *ψ* variables as shown in Equation (12).

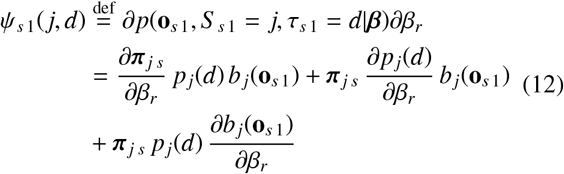

Here *β*_*r*_ can be any coefficient in *β*. Note, that although it is possible for different linear predictors to share some (or all) coefficients (see also Wood et al., 2016), it will usually not be the case that both emission densities and state duration probabilities depend on the same coefficient *β*_*r*_. Hence, it will typically not be necessary to evaluate all three sums in Equation (12) for any given *β*_*r*_. Instead, it will commonly be the case that for any given *β*_*r*_ all but one of the sums evaluate to zero. Note, that similar simplifications will typically be possible when computing the general definition of the recursive update to *ψ* for *T*_*s*_ ≥*t* ≥2, shown in Equation (13). Generally,

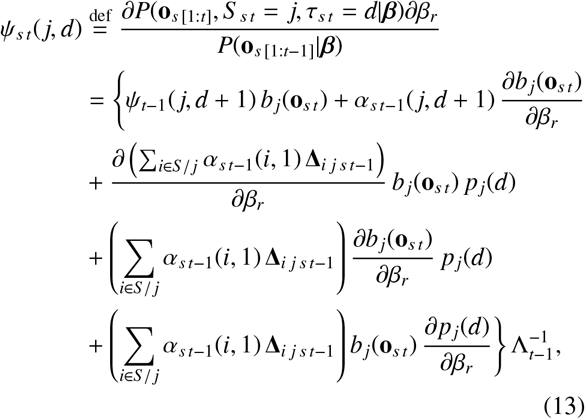

Where

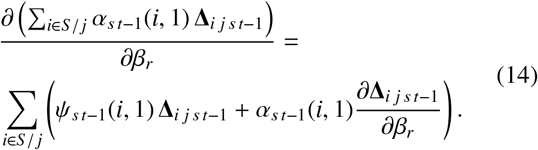

Based on the definitions in Equations (12) and (13) it is clear that

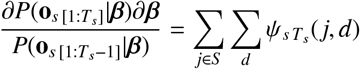

and thus,

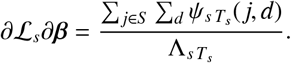

Finally, relying again on the mutual independence of different time-series given *β*,

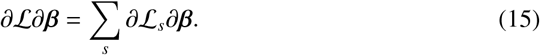

Computing the partial derivatives in Equations (12), (13), and (14) requires partial derivatives of the different densities/probabilities parameterized through different linear predictors. As discussed by Wood et al. (2016), these can typically be evaluated by means of transformation. Appendix B outlines how this can be achieved for the specific derivatives required here. Additionally, note that Equations (12) and (13) can again be computed in parallel for different series *s*, just like the forward variables. Similarly, it is not actually necessary to store *ψ*_*s t*_(*j, d*) for all time-points *t*. As becomes evident from inspecting Equations (12) and (13), computing *ψ* for any given time-point *t* only requires access to the *ψ* variables from the previous time-point *t* −1. Thus, only the *ψ* variables of the previous time-point *t* −1 need to be kept in memory when evaluating the forward recursion.

The information provided so far is sufficient to obtain point-estimates 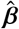 and 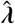 for SMSMs via the L-qEFS update by Krause et al. (2025). The empirical approach, to optimize the Laplace approximate criterion ℒ (*λ*), will likely work well for moderately complex models involving a modest number of smooth functions *f* and random effects. For more complex models, opting for fully Bayesian inference might be preferable instead. For once, the Laplace approximation assumes that, in the vicinity of 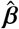, the log-likelihood is approximately quadratic (i.e., that the third and fourth order derivatives are negligible compared to the elements in **H**; see Wood et al., 2016). Additionally, it is necessary that the maximizer of the penalized log-likelihood, 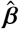, can reliably be obtained for any *λ* – otherwise the Laplace approximation will not hold and updates to *λ* might simply fail. Unfortunately, the surface defined by ℒ (*β*) will typically be all but smooth for complex SMSMs and while ℒ_*λ*_ will typically be slightly easier to navigate, estimation of 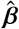 might still get stuck in a local maximum (see Michelot, 2025, for similar discussions).

Theoretically, it is possible to mitigate this risk by restarting the quasi-Newton optimization routine for 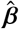, described in more detail by Krause et al. (2025), multiple times from different starting values. Alternatively, a stochastic global optimization algorithm (e.g., simulated annealing; Kirkpatrick et al., 1983) could be used before (or alternated with) the quasi-Newton routine to optimize 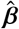. However, for more complex models the associated computational over-head will often be comparable to fully Bayesian inference as discussed in the upcoming paragraphs.

*Fully Bayesian inference* requires the ability to sample from the posterior *p*(*β, λ* | **O**) (see Gelman et al., 2013, for a detailed introduction to Bayesian inference). Since ∂ *ℒ* ∂*β* is available, a No-U-Turn sampler could be used to complete this task efficiently (NUTS; Hoffman & Gelman, 2014). For the NUTS sampler to work, we require the gradient of the joint log-likelihood

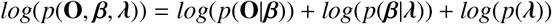

with respect to [*β, λ*] ^⊤^. Here *p*(*λ*) denotes an additional prior distribution placed on *λ* or *ρ* – common choices are wide Gamma and bounded Uniform priors respectively (e.g., Wood, 2020). The partial derivatives of *log*(*p*(**O**, *β, λ*)) with respect to *β* are simply the elements in the gradient ∂ ℒ _*λ*_(*β*)∂*β* of the penalized log-likelihood ℒ_*λ*_(*β*), which can be computed given ∂ ℒ ∂*β* (see for example Wood et al., 2016, for the derivations). Similarly, the partial derivatives of *log*(*p*(**O**, *β, λ*)) with respect to *λ* are the elements in the gradient of *log*(*p*(*β λ*)) + *log*(*p*(*λ*)) with respect to *λ*. The latter is straightforward to compute, considering that the density of *p*(*β λ*) is multivariate normal and the aforementioned choice of standard prior distributions (see Wood, 2017; Wood et al., 2016).

Given these quantities, the NUTS sampler can be used to produce samples *β*^*^, and *λ*^*^ from *p*(*β, λ* **O**). These can be used to derive credible intervals, maximum a posterior (MAP) estimates 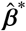, and 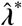, and posterior mean estimates for all quantities involved in the model (i.e., for the coefficients but also individual parameters; see Gelman et al., 2013; Michelot, 2025, for more detailed introductions). For very large models, the requirement to compute ∂ ∂*β* repeatedly in order to generate samples from the posterior might however still become prohibitive.

For very complex models, the alternate sampling strategy proposed by Dewar et al. (2012) will thus be more appropriate. The authors show how to sample from the posterior distribution *p*(*β, λ*, **Z, D**|**O**) by alternating between sampling **Z**^*^, **D**^*^ from *p*(**Z, D**|**O**, *β*) followed by sampling *β*^*^, *λ*^*^ given **Z**^*^, **D**^*^ from *p*(*β, λ* | **O, Z, D**). Here **Z** and **D** are random variables corresponding to the concatenated state sequences and durations spent in all states across all time-series respectively (e.g., Dewar et al., 2012). This approach thus also automatically produces samples from the posterior of state sequences and state durations, which can be useful for further analysis.

Sampling **Z**^*^ and **D**^*^ for the specific EDHMM-like model considered here only requires computing the forward variables. We refer to Dewar et al. (2012) for a discussion of this step and instead focus on the problem of sampling *β*^*^, *λ*^*^ given **Z**^*^, **D**^*^. Conveniently, assuming that no coefficients are shared between different linear predictors, this sampling step can be performed separately for subsets of *β* and *λ*. For example, let 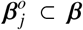 again denotes the sub-set of coefficients involved in the emission model of state *j* (i.e., required by ℱ^*o*^) and let 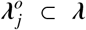 denote the sub-set of regularization parameters involved in any priors place on a subset of 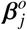. Samples 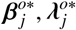 can be drawn from 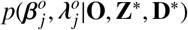. Conveniently, if ℱ ^*o*^ is a member of the exponential family of distributions, the corresponding joint likelihood 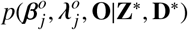 coincides with the one of a Generalized Additive Mixed Model (GAMM) of Location, Scale, and Shape (GAMMLSS, if linear predictors are specified for multiple parameters of ℱ^*o*^; Rigby & Stasinopoulos, 2005; Wood, 2017; Wood et al., 2016) of the subset of emissions **O** that **Z**^*^ indicates have been generated by state *j* (i.e., the rows in **O** for which **Z**^*^ contains *j*) – with any prior on 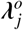 carrying over.

Setting up a NUTS or Gibbs sampler for such a GAMM(LSS) is possible, so that samples 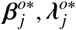 can be generated very efficiently (see Wood, 2016, 2020, for examples). This strategy also generalizes to subsets of coefficients involved in the duration distribution of a given state – the joint likelihood here is of a GAMMLSS of the durations of state *j* in **D**^*^ – and those involved in either the state transition or initial state probabilities – the joint likelihood of coefficients involved in the models of Δ_*i j*_ ∀ *j* ∈ 𝒮*/i* for example, is of a Multinomial GAMMLSS of all transitions away from state *i* in **Z**^*^.

An even more efficient alternative is available if we are content with approximate inference about *β* and *λ*. Specifically, using again the example of 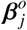 and 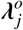, it is also possible to generate samples 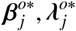 from the large sample Gaussian approximations 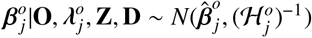 and 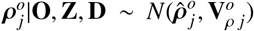, discussed earlier in the context of the Laplace approximation (see also Wood et al., 2016). Here, 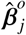 and 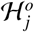 are the maximizer and negative Hessian of the joint log-likelihood 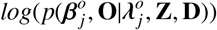. Similarly, 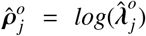 with 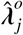 being the maximizer of the corresponding Laplace approximate marginal log-likelihood 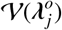. Finally, 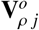 again corresponds to the pseudo-inverse of the negative Hessian of 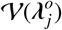 evaluated with respect to 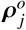 at 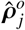 (e.g., Wood et al., 2016). Obtaining these quantities given samples **Z**^*^, **D**^*^ again only requires estimating a GAMM(LSS) of the subset of emissions **O**, generated by state *j* according to **Z**^*^. Then, 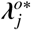 can be sampled from 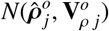 in a first step and 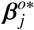 can be sampled from 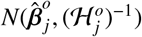 where 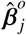 is the maximizer of 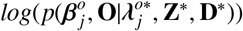 (see Wood, 2017).

This concludes our discussion of the two options, empirical Bayes and fully Bayesian inference, to estimate latent smooth models with a log-likelihood as defined in Equation (9) in the previous section. In section 3 we use the Laplace-approximate approach of the previous paragraph to sample from a multi-level SMSM of the lexical decision data collected previously by Krause et al. (2024). The Supplementary GitHub repository for this paper (see section 3) includes additional examples on how the different estimation strategies presented in this section can be achieved with the mssm Python toolbox (Krause et al., 2025). A critical step of each analysis involving such models, will be to validate whether the model assumptions are reasonable given recorded data. The next section outlines how to achieve this.

#### 2.2.3 Validating Semi-Markov Smooth Models

When working with Generalized Linear Mixed Models (GLMMs) or GAMM(LSS), *model validation* typically involves inspecting the model residuals. Informally, the latter can be considered as the difference between the observed data and the model prediction (e.g., Faraway, 2016; Wood, 2015, 2017). Intuitively, this difference should correspond to random noise if the true data-generating model would be used to generate the prediction. Thus, estimated models producing model residuals that are in clear violation of the expected randomness can be rejected as possible explanations of the data, because they could not possibly have generated it (e.g., Wood, 2015, 2017).

While residuals are commonly used as a tool to validate conventional regression models such as GLMMs or GAMMLSS (see Faraway, 2016; Wood, 2017, for examples), this approach has not yet been adopted widely by researchers using Markov models to recover cognitive processes. This might in part be due to the fact that many different definitions of residuals for HMMs and HsMMs can be found in the literature (e.g., Buckby et al., 2020; Zucchini et al., 2017). Additionally, it is not always clear how these different residual definitions generalize to a multivariate emission process (cf. Kalliovirta, 2008). Thus, we briefly discuss two types of (multivariate) residuals that are applicable to standard EDHMMs and SMSMs (see also Dunn & Smyth, 1996; Kalliovirta, 2008; Langrock et al., 2012; Zucchini et al., 2017). The results section includes practical examples, showing how these residuals can be used to detect problems with these models.

Most studies involving HMMs report some type of pseudo-or quantile-residual (e.g., Buckby et al., 2020; Dunn & Smyth, 1996; Kalliovirta, 2008; Langrock et al., 2012). The specific pseudo-residual considered here, which has also been referred to as *forward* pseudo-residual (Langrock et al., 2012, e.g.,), is defined formally in Equation (16) (see also Dunn & Smyth, 1996). To remain applicable to multivariate emissions, the residual defined in Equation (16) assumes that the *K* signals can be treated as mutually independent, given *β* (e.g., Langrock et al., 2012). As discussed, this assumption is typically justifiable for SMSMs: assuming sufficient random effects are included in the emission models, it is often reasonable to assume that each signal has been generated by a univariate distribution (see also Michelot, 2025). If a multivariate normal distribution is assumed for the emissions instead, then the matrix *N* * *K* matrix **O** first needs to be transformed, based on the estimated covariance matrix for state *j*, so that the *K* columns can again be expected to be mutually independent if the model is correct (see Appendix A).

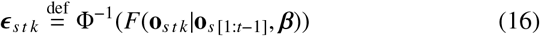

Here Φ^−1^ denotes the quantile function of a standard normal and *F*(**o**_*s t k*_ | **o**_*s* [1:*t*−1]_, *β*) is the cumulative probability of observation **o**_*s t k*_ at time-point *t* for signal *k* and series *s*, given the history of all previous observation vectors **o**_*s* [1:*t*−1]_ for series *s*. For the specific EDHMM-like SMSM considered here, *F*(**o**_*s t k*_ | **o**_*s* [1:*t*−1]_, *β*) can be computed using the forward variables, requiring only a minor modification of the definitions provided in Equations (6) and (7) (see Zucchini et al., 2017). Additionally, if the model is specified correctly *F*(**o**_*s t k*_ | **o**_*s* [1:*t*−1]_, *β*) should be distributed uniformly, so that transforming via Φ^−1^ produces a standard normal variable (see for example Dunn & Smyth, 1996). If the model is specified correctly, the content of each of the *K* vectors *ϵ*_*k*_ should thus look approximately like i.i.d samples from *N*(0, 1). Each vector *ϵ*^*k*^ can thus be inspected like the residual vector of any conventional GLMM and GAMM (see Faraway, 2016; Wood, 2015, 2017, for practical residual plots). A limitation of the residual is that it is typically less sensitive to miss-specified models of the parameters involved in the state sojourn time distributions (see the Supplementary GitHub repository for this paper for a simulation of this tendency).

To address this problem, it will generally be helpful to also consider residuals which utilize an estimate of the latent state sequence **Z**^*^. This includes so-called Viterbi-path residuals, computation of which involves Viterbi decoding to obtain **Z**^*^ (e.g., Buckby et al., 2020). Given **Z**^*^, one can then for example compute deviance or quantile residuals – with the computation depending on the choice for ℱ^*o*^ – of the observations indicated to be in state *j* ∈ 𝒮 (e.g., Dunn & Smyth, 1996; Faraway, 2016; Wood, 2017). Naturally, Viterbi-path residuals are limited in that they are conditioned on the particular choice for **Z**^*^(e.g., Buckby et al., 2020).

However, any residual plots can be generated multiple times for different samples **Z**^*^ obtained either from *p*(**Z** | **O**, *β, λ*) or *p*(*β, λ*, **Z** | **O**).

Given **Z**^*^ and **D**^*^, again denoting an estimate of the duration of all states across all series, residuals can also be defined specifically for properties of the latent process. For example, let 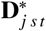 denote the estimated duration of state *j* which started at time-point *t* for series *s*. Based on the definition of the latent process model considered here, we can expect that 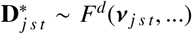 – if the model of the state’s sojourn time is correctly specified. This again enables computation of deviance or quantile residuals for the decoded durations of state *j*. Sampling different **Z**^*^ can again help to alleviate the risk of drawing wrong conclusions based on a particular choice for **Z**^*^ (e.g., Buckby et al., 2020). Consider for example strict left-to-right models: for these models, each state is visited exactly once for every time-series so that only a single residual 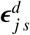 needs to be computed per time-series for state *j*. An average duration residual can then be computed based on samples **D**^*^, which again should approximately be distributed like a standard normal.

In the upcoming Results section we rely on both types of residuals to improve on the model of lexical decision-making proposed originally by Krause et al. (2024).

## 3 Results

In this section we perform a re-analysis of the dataset^6^ collected in the lexical decision experiment presented previously by Krause et al. (2024). Originally, this dataset was analyzed using a standard group-level HsMM approach, here we apply the new framework to be able to account for subject and covariate effects. For a detailed discussion of the experimental setup and covariates used, we refer to the original paper. Here we summarize the aspects most relevant for the reanalysis. The experiment originally featured three different types of stimuli (words, pseudo-words, and random strings) and a continuous measure of each stimulus’ word-likeness, based on Google search result counts. While pseudo-words and random strings are, by definition, non-words, this does not imply that they are all equally word-like (or rather unlike): some pseudo-words might look like miss-spellings of common words while others might feel entirely unfamiliar. Google search result counts act as a continuous proxy measure of these differences in word-likeness and, in contrast to conventional word-frequency measures, can easily be obtained for both words and non-words (see Hendrix & Sun, 2020; Krause et al., 2024, for a more detailed discussion of this measure).

Pre-processing involved low-pass (40 Hz) and high-pass (0.5 Hz) filtering, visual artifact rejection, an independent component analysis to remove eye-related artifacts, and interpolation of missing channels. Afterwards, data was available from 10,948 unique trials (5,402 word, 2,668 pseudo-word, and 2,878 random string trials) collected from 24 subjects. The EEG data were then down-sampled to 100 Hz, base-line corrected by subtracting the mean amplitude in the 100 ms preceding stimulus onset, and epoched so that individual time-series last from stimulus to response onset. Krause et al. (2024) then performed a principal component analysis on the different electrodes and retained the 10 components contributing the most variance (> 90% variance explained), which were normalized per participant. These pre-processing steps are typical for analyses involving the EDHMM model architecture proposed by Anderson et al. (2016), which Krause et al. (2024) used for their analysis as well (see Berberyan et al., 2021; Portoles et al., 2022; van Maanen et al., 2021, for other examples).

Following their approach, the current EDHMM assumes that cognitive processes proceed sequentially from the first to the last process, without the possibility to skip a state. That is, the model is a strict left-to-right EDHMM with a supra-diagonal state transition matrix Δ and *π*_1_ = 1. Thus, the model only requires estimating parameters involved in the state sojourn time distributions and emission models.

The emission model Krause et al. (2024) used was originally proposed by Anderson et al. (2016), and assumes that the EEG process can be characterized sufficiently by *flat* states – periods of variable duration during which the EEG electrodes (or PCA components) have zero mean amplitude and are mutually independent – and *bump* states – microstates of fixed duration during which the EEG is dominated by a stable topography visible across electrodes (or components) (see also Weindel et al., 2024). A more detailed description is provided in Appendix A, alongside the necessary information to compute the density for observed PCA scores during flat and bump states. Notably, state sojourn time distributions only need to be defined for flat states, since bump states are of fixed duration. Krause et al. (2024) used Gamma distributions with state-specific mean parameters but fixed scale^7^ parameters of 0.5 (cf. Anderson et al., 2016; Borst & Anderson, 2015).

Given the number *M* of flat states, estimating the model thus only requires estimating *M* scale parameters and *K* * (*M ™* 1) weights^8^ *µ*_*j k*_, corresponding to the expected value of component *k* during bump state *j*. Krause et al. (2024) used leave-one-subject-out cross-validation to determine the optimal number *M*, which revealed that a model with six flat-states *M* = 6 generalized best to held-out data from all word types (see also Berberyan et al., 2021; Mahé et al., 2015). Notably, they also found that the duration of the penultimate flat state, assumed to reflect the actual decision process, and multiple topographies differed between word types.

### 3.1 Replicating the Six Flat-State Group-level Model

In a first step, we re-estimated their six flat-state model. Because Krause et al. (2024) revealed that the topographies preceding flat states four and five differed between word-types, we define the *µ*_*j k*_ for bump states three and four as shown in Equation 17.

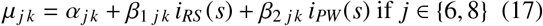

Here *α*_*j k*_ is a state- and PCA component-specific intercept. *i*_*RS*_ (*s*) and *i*_*PW*_ (*s*) are indicator variables that evaluate to 1, if the stimulus of series *s* was a random string and pseudo-word respectively, else 0. *β*_1 *j k*_ and *β*_2 *j k*_ thus correspond to state-, signal- and word-type-specific intercept adjustments, allowing for the topographies of bump states 3 and 4 (*overall* states *j* = 6 and *j* = 8) to vary between word-types. For the remaining bump states, Krause et al. (2024) used the same topography for all word-types. The *µ*_*j k*_ for bump states one, two and five are thus defined only in terms of a state-, and signal-specific intercept *α*_*j k*_ as shown in Equation 18.

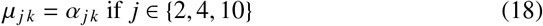

The models for the mean parameters ν _*j*_ of the state so-journ time distributions are similar, but specified for *log*(ν_*j*_) rather than ν _*j*_ directly. Krause et al. (2024) showed that the expected duration of flat states four and five varied between word-types. The models of the (log of the) mean parameters ν _*j*_ for these flat states thus also included state- and word-type-specific intercept adjustments, while the models for the remaining states again included only a state-specific intercept. Note, that the *µ*_*j k*_ (and ν _*j*_) lack time-series *s* and time-point *t* subscripts, because the original model by Krause et al. (2024) only includes group-level effects. Later in this section we present a more appropriate hierarchical model alternative.

To estimate the resulting model, the quasi-Newton approach utilized by the L-qEFS update (e.g., Krause et al., 2025) was used to maximize the log-likelihood as defined in the Methods section. As expected, the resulting model produces estimates virtually indistinguishable from those obtained by Krause et al. (2024), which are shown in Figure 2.

**Figure 2.**
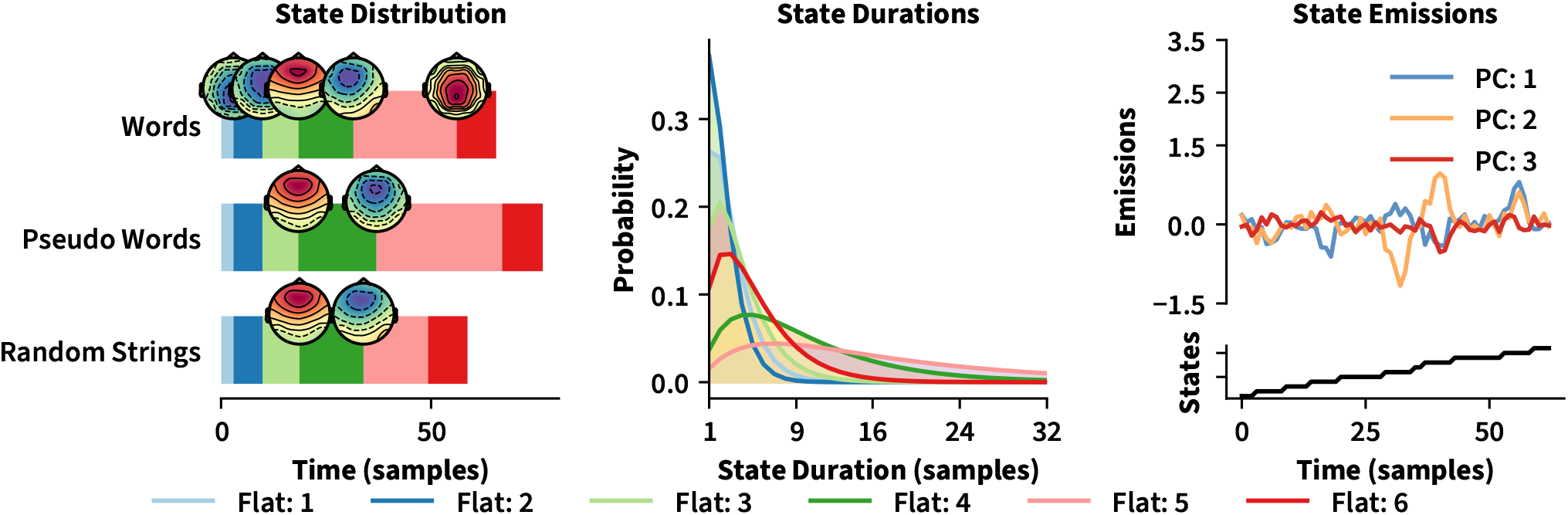
Overview Six Flat-State Model. Figure 2 shows an overview of the re-estimation of the six flat-state model of lexical decisions proposed originally by Krause et al. (2024). The left-most panel shows the average duration of each flat state separately for every word-type. These were estimated from the Viterbi-decoded state sequences. The plot also contains the estimated bump topographies, preceding the onset of each flat state. The central panel shows the average (over word-types) probability mass function for the duration of each flat state – note how later flat states are generally associated with wider distributions, in particular the penultimate flat state. The right-most panel visualizes the emission and latent process assumed by the model: during a flat state the signal is noisy with an expected amplitude of zero. Entering bump states results in a weighted half-sine being added to the noisy process. The peaks of the weighted half-sines from all electrodes form the topographies shown in the left-most panel.

Notably, four different topographies can be distinguished for words in the left-most panel of Figure 2: two topographies featuring central-posterior negativity, a third topography resembling the mid/frontal old-new (N170) ERP component, a fourth topography featuring central-anterior negativity, and a final, fifth, topography featuring parietal-posterior positivity (see the top bar of the bar-plot). Previous work suggests similar patterns to be related to the onset of visual processing, orthographic familiarity assessment, decision-making, and response initialization respectively. The duration of the fifth flat generally dominated the overall reaction times for all word types (see central panel of Figure 2 for average duration mass functions). However, the duration of this flat state also varied strongly between word types (see the left-most panel of Figure 2), a pattern that previous models suggest is typical for (lexical) decision-making (e.g., Ratcliff et al., 2004; Wagenmakers et al., 2008).

In a second step we computed forward pseudo-residuals for this six stage model, as defined in the methods section. As discussed in section 2.2.3, pseudo-residuals are typically computed for each of the *K* multivariate signals (PCA components in this case). Because it is difficult to show plots for all 10 PCA components, we only show the plots for the first component, with the understanding that the plots for the other components show similar patterns unless otherwise noted. Additionally, the Supplementary GitHub repository for this paper contains code to create the plots for all components. Figure 3 shows quantile-quantile (QQ-plot), time-course, and ordered residual plots as well as an estimate of the auto-correlation function of the residuals for different time-lags. While the residuals appear to be approximately normally distributed marginally (see the QQ-plot; other PCA components show stronger deviations), the variance of the residuals appears to change over time (second plot from the left) and to differ between subjects (third plot from the left). Similarly, the residuals continue to show strong temporal correlations, even for distant time-lags (right-most plot). As such, there appear to be remaining dependencies in the data that the model cannot account for.

**Figure 3.**
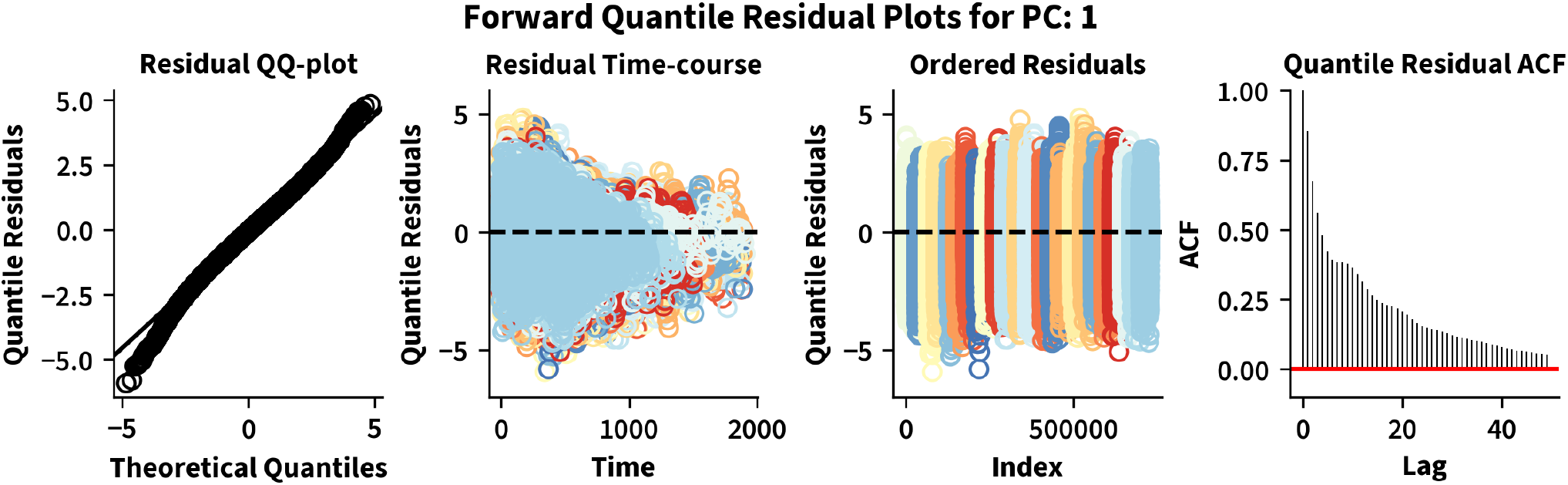
Residuals of Six Flat-State Model. Figure 3 shows different plots of the forward pseudo-residuals computed for the six flat-state model of lexical decisions shown in Figure 2. The left-most panel plots the observed residual quantiles against the theoretical (standard normal) quantiles to assess deviations from normality. The second plot from the left plots the residuals against time, with residuals from different subjects being colored differently. Note, that the variance in the residuals appears to decrease over time. The third plot from the left shows the residuals in the order of the data, again colored by subject. Note, that there appears to be a difference in the variance of the residuals between subjects. The final plot shows the estimated auto-correlation function of the residuals for different time-lags. Evidently, residuals of this model remain highly correlated.

A possible explanation for the remaining temporal correlation of the residuals is the comparatively simple emission model used here (see the right-most panel of Figure 2 for a visualization of the model’s prediction for the emissions for the first three PCA components): instead of assuming that emission deviations from the expected value (zero during flat states, topography-dependent during bump state) are i.i.d realizations of a standard normal, it might be more appropriate to assume that these deviations are temporally correlated. The strong temporal correlations visible in Figure 3 persist over multiple lags, which suggests that a multi-lag auto-correlation model of the deviations might be required. However, because such a model would be very expensive to compute, we considered an auto-correlation model of lag 1 (i.e., an ar-1 model) instead – the details of this model are provided in Appendix A. Based on a visual inspection of the auto-correlation function in Figure 3, we chose *ρ* = 0.7 for the value of the auto-correlation weight.

Figure 4 shows the updated residuals for the six flat-state model including an ar-1 model of the emission deviations. The auto-correlation function has improved substantially, although weak temporal correlations between shorter time-lags remain visible (see right-most panel of Figure 4). Residuals also appear to deviate slightly more from marginal normality, and their variance continues to decrease with time and to differ between different subjects. This might in part be a result of an overly simplistic model of the state sojourn time distributions: In a follow-up analysis Krause et al. (2024) showed the expected duration of all flat states to vary as a word-type-specific function of word-likeness. Additionally, the expected duration of flat states varied smoothly over trials – and did so differently for different subjects. The current model of the state sojourn times, a Gamma distribution with state-specific mean parameter, cannot account for either of these patterns.

**Figure 4.**
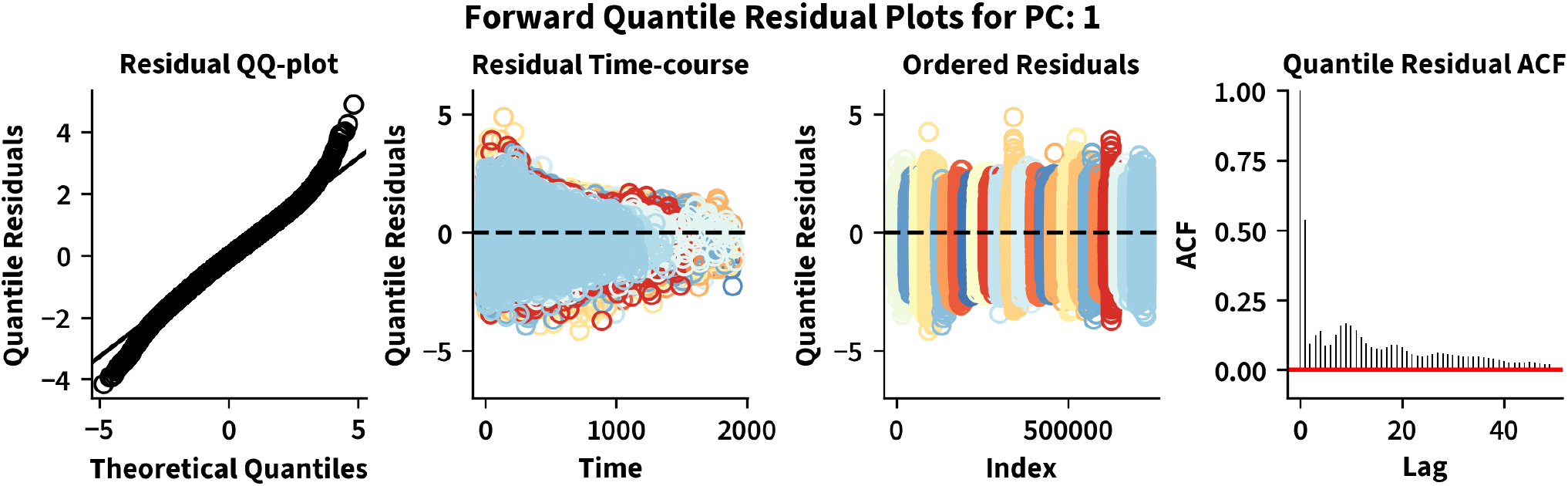
Residuals of Six Flat-State Model Including an ar-1 Model of the Emissions. Like Figure 3, Figure 4 shows different plots of the forward pseudo-residuals, but computed for the the six flat-state model including an ar-1 model of the emission deviations. Notably, while the residuals of this model appear to deviate slightly more from marginal normality (see left-most plot) temporal correlations between residuals are substantially weaker (see right-most plot). The variance of the residuals still appears to decrease over time and to differ between subjects however (see central plots).

To test for the correctness of the model of the state sojourn times, we computed average residuals of the duration of each flat state for every time-series, as discussed in the final part of section 2.2.3. The average duration residuals were computed based on 100 samples obtained from the posterior of state sequences given observed data and maximum likelihood estimates of all model parameters. Different visualizations of these residuals are depicted in Figure 5 for flat states 3-6 (flat states 1 & 2 showed similar patterns as flat state 3 and were thus omitted from the Figure to improve legibility).

**Figure 5.**
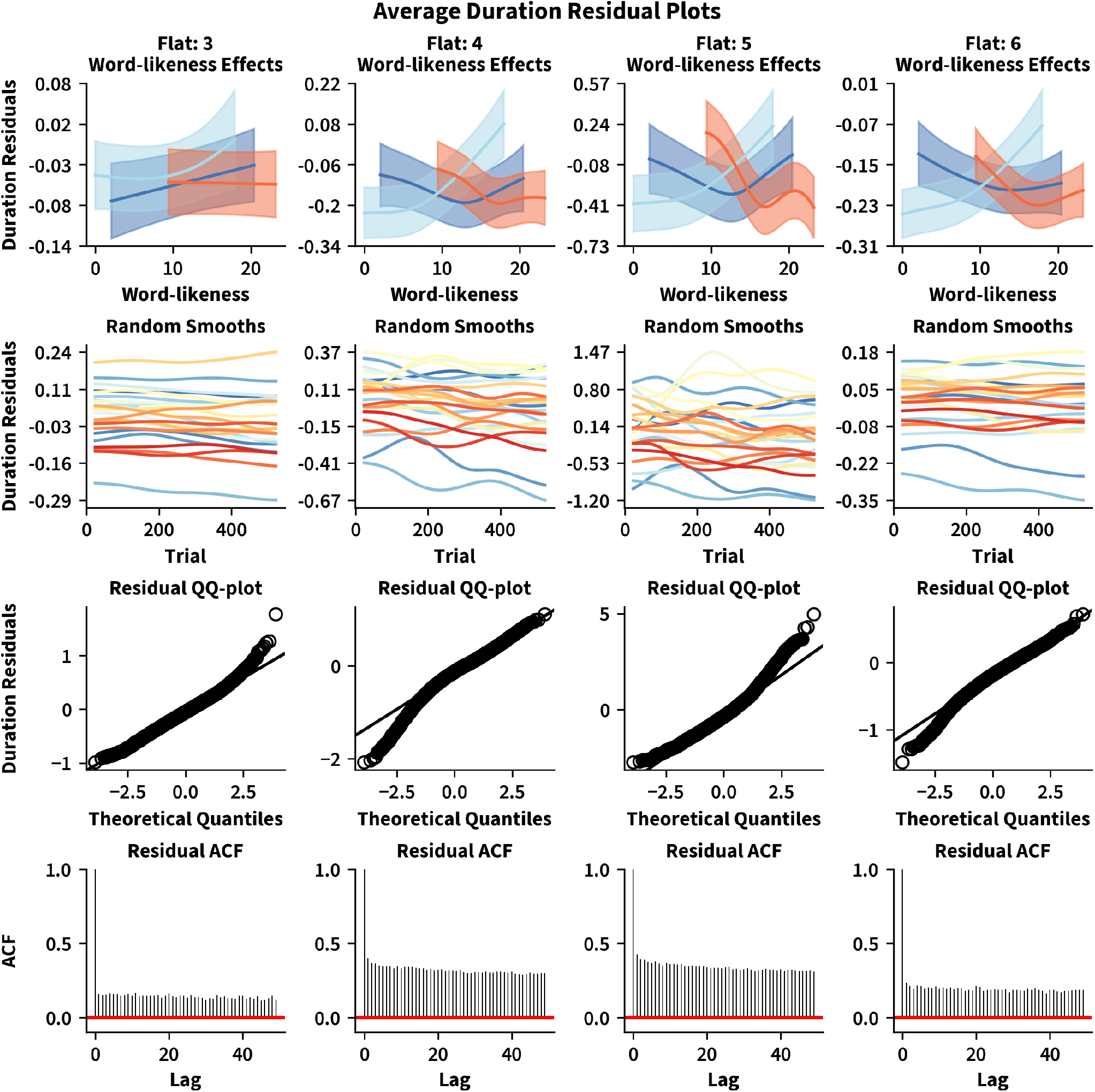
Average Flat-State Duration Residuals. Figure 5 shows plots of the average duration residuals for all flat states, computed for the the six flat-state model including an ar-1 model of the emission deviations. The first row shows, separately for each word-type, the estimated effect of word-likeness on the expected residual. As observed previously by Krause et al. (2024), later flat states show more complex effects of word-likeness (note the y-axis differences). The second row shows estimates of the subject-specific change in the expected residual over trials. The deviations also appear to be greater for later flat states – in particular the penultimate flat state. The last two rows depict QQ-plots and estimates of the auto-correlation function of the residuals: the average duration residuals appear to deviate slightly from normality and generally show strong temporal correlations across different lags (i.e., trials). This suggests that expected flat durations vary between subjects and over the time-course of the experiment.

The first row of Figure 5 visualize estimates of the state- and word-type specific effects of word-likeness on the expected duration residual. Similarly, the second row shows subject-specific changes in the expected residual over the time-course of the experiment. These were obtained by fitting additive mixed models (e.g., Wood, 2017) to the state-specific average duration residuals. If the model of the state sojourn times were correct, the expected value of the duration residuals should be zero and constant, i.e., insensitive to changes in word-likeness or time spent in the experiment. Instead, the clear deviations from this pattern, visible in the top rows of Figure 5, suggest that the expected duration of the different flat-states is not constant, but rather an additive combination of state-, subject-, and word-type-specific smooth non-linear functions of word-likeness and the experimental time-course. The fact that the state-specific duration residuals deviate from normality and show strong temporal correlations across multiple trials further suggests that the current model of the sojourn time distributions is too simple (see last two rows in Figure 5).

As mentioned, the fact that the current model of the state sojourn times does not account for any subject-specific differences, could partially explain why the variance of the pseudo-residuals, shown in Figure 4, appears to vary between subjects. It is more likely however, that the emission process model also needs to account for subject-specific differences. The current model assumes that the state topographies preceding flat states are the same for all subjects (see Equations 17 and 18). Even if all subjects complete the same sequence of processes however, the topographies marking the onset of a new process could very well be different for different subjects. In fact, it might not even be reasonable to assume that all subjects complete the same sequence of processes (on all trials): even for a simple task as lexical decision-making, different subjects might employ slightly different strategies.

### 3.2 A Hierarchical Six Flat-State Model

To account for these possibilities, a multi-level or hierarchical latent semi-Markov smooth model is required with additive mixed models for the *µ* and ν parameters. For the *µ* parameters we keep the group-level structure the same as before and only add subject-specific random effects. Therefore we again need to define different effect structures for bump states three and four on the one hand and bump states one, two, and six on the other. The updated models are shown in Equations 19 and 20.

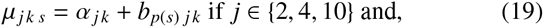

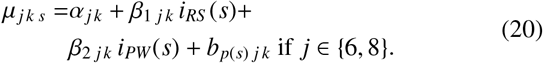

Here, *b*_*p*(*s*) *j k*_ denotes a state- and PCA component-specific random intercept (adjustment) for subject *p*(*s*) from which series *s* was recorded. The inclusion of the *b*_*p*(*s*) *j k*_ allows for different topographies to be estimated for each subject. Additionally, we again need to define mixed models for the (log of the) mean of the state sojourn time distribution ν _*j s*_, which, as shown in Equation (21), can now differ for series *s* and flat state *j*.

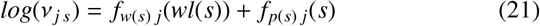

Here, *wl*(*s*) defines the word-likeness of the stimulus present for series *s*. Correspondingly, *f*_*w*(*s*)_(*wl*(*s*)) denotes a state- and word-type (*w*(*s*) returns the word-type of the stimulus present for series *s*) specific smooth function (including an offset) of the word-likeness *wl*(*s*) of the stimulus present for series *s*. Finally, *f*_*p*(*s*) *j*_(*s*) is a state- and subject-specific *random smooth function* (i.e., subjected to a proper full-rank prior) of the experimental time-course (i.e., *s*; see Wood, 2017).

The resulting latent smooth model is quite complex, requiring estimation of 2910 coefficients *β* and 80 inverted variance/regularization parameters *λ*. Therefore, we relied on the Laplace-approximate Bayesian estimation approach discussed in the methods section, alternating between sampling state and state sojourn time sequences **Z**^*^, **D**^*^ and using normal approximations to sample *λ* | **O, Z**^*^, **D**^*^ and *β* | **O**, *λ*, **Z**^*^, **D**^*^ (see also Dewar et al., 2012).

Two chains were sampled for 1250 iterations, with the first half being subsequently discarded and only the second half of each chain being used for analysis. Trace plots suggested that the marginal posteriors for some random intercepts (mostly those estimated for subjects 1 and 18) and *λ* values might be multi-modal, with both chains converging to different but stationary modes. To test for the possibility of additional modes, both chains were re-initialized based on averages taken over different sampling windows of the original chains and sampling was continued for 50 additional iterations: for coefficients showing multi-modality, both chains virtually immediately jumped back to one of the previous modes. For the remaining coefficients, visual inspection of the trace plots suggested that the original chains converged approximately to the same mode. However, 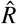 remained (slightly) above 1.1 for some of these coefficients, suggesting that sampling would have benefited from additional iterations to really establish approximate convergence (see Gelman et al., 2013, for an introduction to convergence statistics and a more detailed discussion).

Figure 6 shows an overview for this hierarchical six flatstate latent smooth model. The left-most panel shows posterior means of the group-level bump topographies and flat state duration estimates. While the group-level estimates of the bump topographies are comparable to those obtained with the simpler model (visualized in Figure 2), the estimated durations of the flat states show greater differences compared to the simpler model. Similarly, for some participants the subject-level estimates of flat state durations (see central panel in Figure 6) and topographies alike (see rightmost panel in Figure 6 and Figure C1 in Appendix C1) deviate substantially from the group-level estimates obtained from the simpler model.

**Figure 6.**
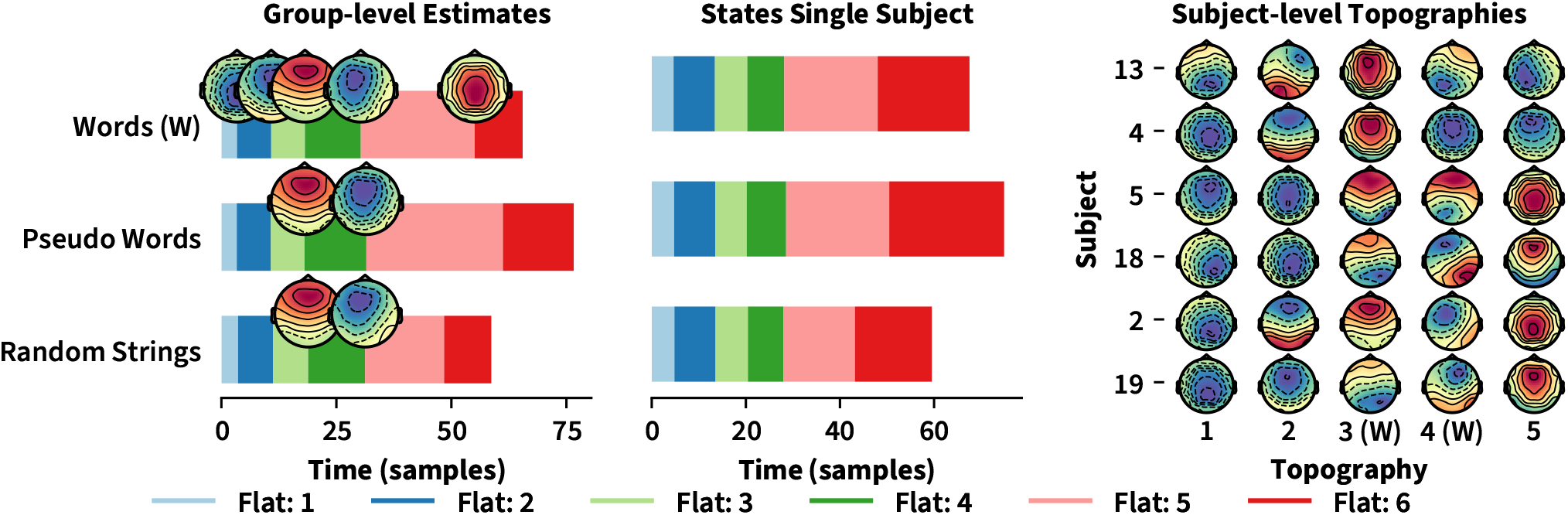
Six Flat-State Multi-level Semi-Markov Smooth Model. Figure 6 shows an overview of the six flat-state multi-level semi-Markov smooth model of lexical decisions. The left-most plot matches the left-most plot in Figure 2 and shows the estimated duration of each flat state, as well as the estimated topographies preceding these states. Importantly, the topography and state duration estimates were obtained using only the posterior mean estimates for the *group-level coefficients* (i.e., *α*_*j k*_, *β*_1 *j k*_, *β*_2 *j k*_, and the coefficients involved in the *f*_*w*(*s*) *j*_(*wl*(*s*))) in the models of *µ* and ν defined in Equations (19-21). In contrast, the central plot shows the state duration estimate obtained for a particular subject, based on the posterior mean estimates for *all* coefficients (i.e., without zeroing random coefficients). Similarly, the right-most plot shows a matrix of the posterior mean estimates of the subject-specific bump topographies for words (“W”) and a subset of select subjects (see Figure C1 in Appendix C for a version including all subjects). Note, that while some subjects show bump topographies similar to the group-level estimates in the left-most panel, other subjects show bump topographies that differ systematically from the group-level estimates – which is particularly evident for the bump topography preceding the fifth flat (i.e., topography 4, here only shown for words).

The differences caution against over-interpreting decoding estimates from a model including only group-level effects: in such a model the decoding is forced to determine state estimates based on topographies/emission model parameters that might not actually be optimal for a particular subject. As a result, decoding estimates from such a model might exaggerate differences between states, suggesting that distinct states might reflect distinct processes.

In contrast, the subject-level variations in the estimated topographies suggest that flat-states four and five might actually reflect a mixture of partially overlapping processes or even the evolution of a single (decision) process over time, rather than distinct processes: for a sub-group of subjects (e.g., subjects 5 and, to a lesser degree, 13 in Figure 6 as well as subjects 3, 8, 15, 21, 23, and 24 in Figure C1 included in Appendix C), the subject-level estimates (also based on posterior means) of the fourth topography actually show strong positivity, rather than negativity. These estimates thus remain qualitatively more similar to the group-level estimate of the third topography, again resembling the familiarity-related N170 ERP component (e.g., Krause et al., 2024). A possible explanation: topographies three and four are snapshots of an evolving decision process, with some some subjects (e.g., 5 and 13 in Figure 6) relying more than others on evidence accumulated early during this process. This could suggest that a sub-group of participants might have employed a different strategy than the rest of the group, to reach the final decision.

Further evidence for the aforementioned hypothesis stems from the posterior mean estimates of the *f*_*w*(*s*) *j*_(*wl*(*s*)), the state- and word-type-specific smooth functions of word-likeness included in the models of the expected duration of state sojourn times. The corresponding estimates are visualized in Figure 7, and differ slightly from the estimates obtained in a follow-up analysis by Krause et al. (2024). Specifically, the posterior mean estimates computed here mainly show differences between the first three and the last three flat states. As observed by Krause et al. (2024), the effect of word-likeness is generally more attenuated in these first three flat states. However, in contrast to their previous findings there appear to be no differences between word-types (see Appendix D for credible intervals of the difference between word-types) in these first three flat states. Additionally, the estimated effects of word-likeness (and word-type) in the last flat states are generally more similar across states – particularly for flat states five and six. This also explains why both flat states five and six now show similar main effects of word-type on the average duration (see pink and red bars in left-most panel in Figure 6). While Krause et al. (2024) also observed similar effects of word-likeness in flat states five and six, they generally observed more pronounced differences between flat states – particularly between flat states four and five.

**Figure 7.**
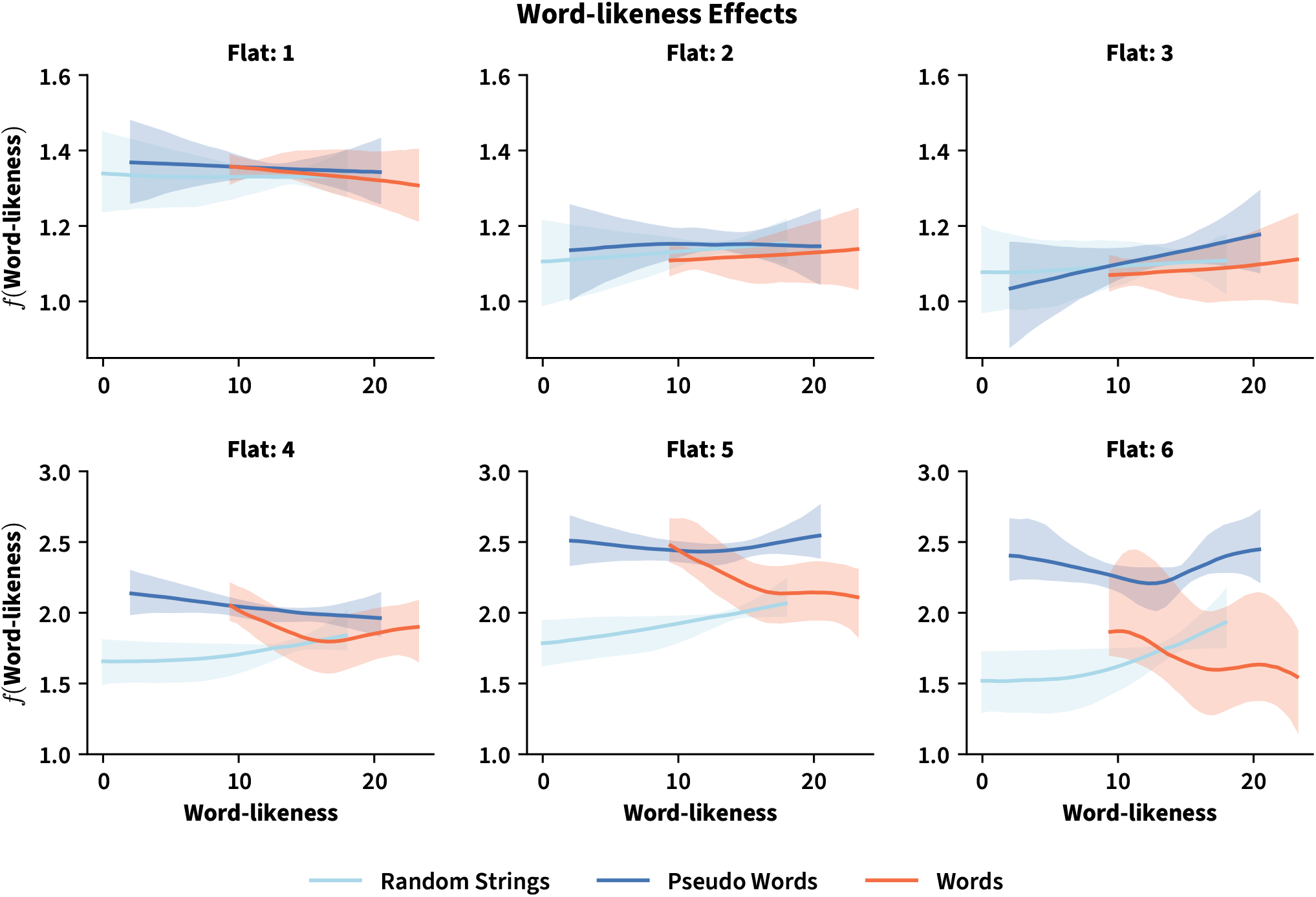
Word-likeness Estimates from Multi-level Semi-Markov Smooth Model. Figure 7 shows an overview of the posterior mean estimates of the smooth functions *f*_*w*(*s*) *j*_(*wl*(*s*)) involved in the models of the mean parameter ν_*j*_ of the state sojourn times. Each panel corresponds to a different flat state, and visualizes the estimated smooth for all three word-types. Shaded areas are 95% posterior credible intervals for the shape of the word-likeness smooth.

A possible explanation for these differences is the fact that the results for the multi-level model are based on posterior mean estimates – the MAP estimates might produce estimates more comparable to those reported by Krause et al. (2024). However, the fact that word-likeness effects are more similar in the last three flat states also supports the aforementioned hypothesis that some or all of these flats might actually reflect a continuously evolving process, rather than distinct cognitive processes.

More work will be necessary to establish this, since even the complex multi-level model presented here neglects an important source of subject-level differences: Specifically, for some subjects (e.g., 4 and 13 in Figure 6 as well as 1, 7, 12, and 14 in Figure C1 included in Appendix C), the final topography, preceding the final flat state, continues to show strong negativity and thus remains more similar to the word-type-specific group-level estimates of the fourth topographies preceding the fifth flat. This observation aligns with the findings by Krause et al. (2024), who showed that an additional sub-state might be visible between flat states five and six. The authors suggested that this sub-state might only be necessary for pseudo-words, reflecting a more complicated decision process for these types of non-words. The analysis performed here suggests, that this extra sub-state might also be present only for a subset of subjects. This could imply that this sub-group of subjects dedicated additional processing time to the decision process or that these subjects relied on additional information, not considered by other subjects.

Importantly, the multi-level model presented here did not account for this sub-state. As a result, parts of the duration of the fifth flat state might have been (incorrectly) attributed to the duration of flat states four and/or six. Specifically, we would expect that, when including an extra topography for selected subjects for which the final topography currently shows strong negativity, their estimates of the final topography would again become more similar to the current group-level estimate showing parietal positivity (see left-most panel in Figure 6). In consequence, the estimated effects of word-likeness on the duration of the different flat states might changes as well and potentially become more similar again to the effects observed by Krause et al. (2024).

Additionally, while the average residuals of flat state durations have improved considerably for this multi-level model (see Appendix E), the forward pseudo-residuals improved only slightly (see Appendix E), with some subjects continuing to show greater variance in the residuals. Exploring an extra sub-state (i.e., topography plus flat state) for the aforementioned individuals might be sufficient to address this, but it might also be necessary to consider a more complex random-effect structure for the models of the *µ* in general.

To summarize, the results presented here generally support previous findings, which suggested that five prototypical EEG topographies precede the response in lexical decision tasks (e.g., Krause et al., 2024). Rather than reflecting distinct processes, the topographies preceding flat states four and five might reflect the continuous evolution of a single decision process over time. Additionally, even for a simple lexical decision task there appear to be systematic strategic differences between subjects: some appear to rely more strongly, or even exclusively, on evidence available early during the decision process, while others seem to dedicate extra processing time to reach an accurate decision. Extensions of the model presented here remain necessary, but are outside the scope of the current paper, as the main goal was to illustrate the new framework.

While the modeling framework presented here is sufficiently general to accommodate even complex future additions, such as subject-specific extra states, practical application is complicated by the considerable computing resources required to estimate these models. This and other open challenges will be discussed in more detail in the upcoming section.

## 4 Discussion

In this paper we discussed hierarchical semi-Markov smooth models (SMSMs) of latent neural states, and how to estimate them. SMSMs improve on existing methods to discover cognitive processes from neuro-physiological recordings because i.i.d random effects and smooth functions of covariates can be included in the models of individual parameters. This allows SMSMs to account for subject-level heterogeneity and potentially non-linear covariate effects in the latent and emission process.

To demonstrate the usefulness of SMSMs, we re-analyzed the lexical decision data presented previously by Krause et al. (2024). The primary objective of this analysis was to give an impression of the kind of research questions that can be investigated with these models; “Do all subjects complete the same processes in the same order?” “Do the expected duration of processes or state transition probabilities change smoothly over the time-course of an experiment or as a function of experimental covariates?” – because they can incorporate random and smooth effects, SMSMs can help answer these and related questions.

Additionally, we wanted to demonstrate that existing HsMM models, even very specific ones like the left-to-right EDHMM presented by Anderson et al. (2016) and used by Krause et al. (2024), can quickly be transformed into SMSMs. Conceptually, the SMSM framework presented here is not a replacement for a specific method or type of model used. Instead, the framework presented here can be applied to extend any kind of HsMM (or HMM; see Michelot, 2025) for which the log-likelihood and gradient can be computed. As such, the framework is agnostic to the specific choice of emission density and whether or not initial state and state transition probabilities are to be estimated or not. Moreover, SMSMs can be applied to univariate and multivariate emission processes alike: cross-temporal dependencies between multivariate signals can either be accounted for by choosing an emission model explicitly accounting for such dependencies (e.g., Vidaurre et al., 2016, 2018) or by including sufficient random effects in the emission models so that the signals can reasonably be treated as realizations from univariate densities (see Michelot, 2025, for similar arguments).

Because the focus was on introducing the SMSM framework, the analysis presented here involved some shortcuts: the six flat-states solution determined by Krause et al. (2024) was used as a starting point and the number of states was subsequently kept fixed. Despite these limitations, which we discuss in more detail in a later part of this section, the analysis performed here provided important theoretical insights, further highlighting the usefulness of the SMSM framework.

First of all, the analysis revealed substantial between-subject differences for some of the *EEG topographies* – essentially the emission process parameters of the specific EDHMM used here (see right-most panel of Figure 6). Previously, Anderson et al. (2016) argued that these topographies reflect the onset of a distinct “processing stage”. However, many of the subject-level estimates of the word-type specific topographies preceding the fifth flat state deviated considerably from the group-level estimates featuring strong central negativity (see left-most panel of Figure 6). For some subjects, the estimates could better be described as a mixture of the topographies preceding the fourth and fifth flat state (see subjects 3, 5, 8, 13, 15, 21, 23, and 24 in the right-most panel of Figure 6 and Figure C1 in Appendix C). For other subjects, the estimated topographies preceding the fifth flat state showed no sign of central negativity at all. As discussed in the Results section, the topographies preceding flats four and five could thus also reflect a time-continuous emission process of a single latent process rather than two distinct processing stages.

The emission process model by Anderson et al. (2016) assumes that the EEG returns to an average amplitude of zero after briefly resembling one of the state-specific topographies, and thus does not account for such a continuous emission process. Instead it might be more appropriate to merge flat states four and five and to model the emissions of this merged state with a multivariate Gaussian with a mean vector varying smoothly as a function of the duration of the state. Theoretically, this only requires minor modifications of the likelihood function defined in the Methods section. Yet another alternative, that does not require any modification of the log-likelihood defined here, would be to rely on a multi-variate auto-correlation model for the emissions of this merged state (e.g., Vidaurre et al., 2016).

Additionally, the analysis revealed that including an extra topography and flat state for some subjects might considerably improve the fit for these subjects (see subjects 1, 4, 7, 12, 13, and 14 Figure 6 and Figure C1 in Appendix C). Specifically, the analysis revealed that, for some subjects, the subject-level estimate of the final topography was less similar to the group-level estimate of the final topography, featuring strong parietal positivity. Instead, the subject-level estimates were more similar to the group-level estimate of the earlier topography, preceding the fifth flat and featuring strong central negativity. This ability, to accommodate subject-specific deviations from the group-level state trajectory, is a major advantage SMSMs have over pure group-level models.

As discussed briefly in the Results section, we hypothesize that the corresponding subjects take more time to reach the actual decision or consider additional information not accessed by most subjects. Hence, we would expect that the final topography of these subjects becomes more similar to the final group-level topography again, when estimating an extra topography and flat state for these subjects. Intriguingly, Krause et al. (2024) suggested that such an extra state might actually be completed by all subjects, but only for pseudo-words. With the SMSM framework presented here it is possible to investigate which of these competing hypotheses, or perhaps a combination, is more likely.

More generally, the subject-level deviations observed here also caution against over-interpreting estimates obtained from models including only group-level effects: state estimates decoded from group-level estimates alone might, for example, exaggerate differences in the duration or emissions of individual states which can generate the impression that these states reflect distinct processes. This reinforces the point raised briefly in the Introduction, that Semi-Markov states can, at best, be considered a reflection of a combination of temporally (partially) overlapping processes (e.g., Baker et al., 2014; Pieramico et al., 2025; Vidaurre et al., 2025).

An important factor that contributed to the decision not to investigate a model with a subject-specific extra state or a different emission process model for a combination of the fourth and fifth flat state, is the considerable computational complexity associated with estimating SMSMs. Obtaining 1250 samples from the final model reported here took more than a week and a dedicated server with 40 CPUs. This drastically complicates the task of selecting between two or more candidate models – even when an estimate of or prior knowledge about the number of states and appropriate emission distribution/model is available so that the candidate models differ only in their effect structure. While previous work typically considers only a single emission distribution/model shared by all states, often motivated by theoretical considerations, it is common to estimate the effect structure and number of states alongside model parameters (e.g., Borst & Anderson, 2015; Vidaurre et al., 2025; Weindel et al., 2024). We return to the problem of determining the number of states in the final part of this discussion and for now focus instead on the problem of determining an appropriate (hierarchical) effect structure for the models of the different parameters given (an estimate of) the number of states (and emission distributions/models).

### 4.1 Determining an Appropriate Effect Structure

This “simpler” problem is already far from trivial, because the space of possible SMSMs is extremely vast – even for simple experiments involving only a few manipulations and covariates. Taking into account the aforementioned high computational complexity of the log-likelihood and gradient functions, a brute-force cross-validation approach over a grid of different effect structures for parameters involved in the emission and latent processes and any remaining hyperparameters (e.g.,the *ρ* value of an ar-1 process) will often be impossible. For that reason, we instead chose the value for the *ρ* parameter of the ar-1 model of emission deviations and the effect-structure of the state sojourn time models based on inspections of residual visualizations and previous work.

To be clear, modifying a model based on patterns visible in the residuals is a valid strategy – not every extension has to be informed by a significance test or a selection procedure. Indeed, the results from the analysis performed here again highlight the importance and usefulness of residual visualizations and model validation in general: the forward pseudo-residuals (see Langrock et al., 2012) of the six flat-state model showed strong temporal dependencies. Adjusting the specific emission model proposed by Anderson et al. (2016) to incorporate an ar-1 model of the PCA deviations (see Appendix A) largely accounted for these temporal dependencies – providing immediate justification for the added complexity of the resulting model.

In line with this, we would recommend to generally rely heavily on model validation for the task of selecting an appropriate (hierarchical) effect structure given (an estimate of) the number of states. Ideally, this search would already start with a reasonably complex effect structure, involving sufficient random effects in the models of all parameters (c.f. Bates et al., 2018), determined based on theoretical considerations. Subsequently, the different types of residuals discussed in the methods section could then be used to check if the data contains substantial remaining dependencies that the model cannot account for. Additionally, a formal selection among a small number of candidate models can be performed without having to complete an expensive cross-validation procedure.

Instead, the selection could be based on any of the available Bayesian selection criteria which can be computed from sampling output (see Gelman et al., 2013, for an overview). Alternatively, a conditional version of the Akaike Information Criterion (AIC) can be computed as 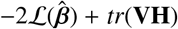, where 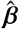 is the maximizer of the penalized log-likelihood given 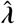 obtained as discussed in section 2.2.2, **H** is the negative Hessian of the log-likelihood (or a quasi-Newton approximation; see Krause et al., 2025) evaluated at 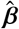, and *tr*(**VH**) denotes the trace of **VH** (Wood et al., 2016). **V** can either be based on the covariance matrix of the Laplace approximation to the posterior *β*|**O**, *λ* or estimated from samples of the posterior *β, λ* | **O** in which case the AIC will automatically be corrected for uncertainty in the estimate 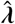 as well (see Wood, 2016; Wood et al., 2016).

### 4.2 Determining the Number of States

A data-driven analysis designed to determine the optimal number of Semi-Markov states alongside an appropriate effect structure faces the same problems outlined before. However, while computational complexity was already a concern for the task of determining an appropriate effect structure, a brute-force cross-validation approach over a grid of different effect structures for parameters involved in the emission and latent processes, number of states, hyper-parameters, and (optionally) different emission and state sojourn time distributions/models becomes intractable.

A potential alternative might be to adopt a sequential selection strategy – not unlike what was done for the analysis presented here: first, cross-validation could be used to determine the optimal number of states, given a fixed and relatively simple group-level effect structure and choices for emission and state sojourn time distributions. The second step, determining an appropriate hierarchical effect structure as well as values for any hyper-parameters could then be addressed as discussed in the previous section, based on the number of states estimated in the first step. For example, Krause et al. (2024) used leave-one-out cross-validation (LOOCV) and only accepted additional states if this reduced the cross-validation error for a majority of subjects. In consequence, it could be expected that the recovered EEG topographies should make for good group-level estimates in a hierarchical model as well. Indeed, the hierarchical model presented here produced very similar group-level estimates (compare the left-most panels of Figures 2 and 6), despite the fact that the hierarchical model also included an ar-1 model of the emission deviations.

Naturally, there are many ways in which this first step might fail to produce a useful estimate for the number of states. For example, if, in truth, word-likeness had a (non-linear) group-level effect on the EEG topographies, LOOCV, based on a model treating the topographies as constant, would be unlikely to result in a useful estimate of the number of states. As such, success of such a sequential selection strategy cannot be guaranteed^9^. Fortunately, the framework presented in this paper, implemented in the mssm Python toolbox (version ≥1.2.0; Krause et al., 2025), allows to thoroughly investigate the usefulness of this and any other strategy to estimate the number of states. Additionally, careful model validation, based on the different residual types considered here, will at least allow to determine whether the final model obtained by any such strategy remains highly unlikely to have generated the collected data. Similarly, inspections of these residuals can also guide the search for an appropriate effect structure.

### 4.3 Conclusion

In summary, the SMSM framework offers a valuable addition to the toolkit of researchers wanting to discover cognitive processes from neuro-physiological recordings. SMSMs can account for subject-level heterogeneity in the latent and emission process and can include smooth functions of experimental covariates in the (mixed) models of individual parameters. Because the computational complexity of estimating (or sampling from) SMSMs is considerable, more research is necessary to validate and determine effective data-driven model selection strategies. To facilitate such research, all the algorithms considered in this paper have been implemented in the mssm Python toolbox.

## Appendix A

### Working with Emission Densities Conditional on Residual State Duration

As discussed in the main paper, Anderson et al. (2016) set out to define a more appropriate model of EEG emissions, based on the observation that several theories predict that the EEG signal can be described as a multivariate oscillatory noise process with zero expectation most of the time. Only in a brief period following a shift in processing, assumed to coincide with the onset of a new cognitive process, is the signal believed to deviate systematically from this process.

Specifically, the onset of a new process is assumed to result in the addition of a weighted half-sine of fixed duration to the EEG signal at each electrode. Anderson et al. (2016) thus suggest to distinguish between the emissions generated while in a “flat state” and the emissions generated during these “bumps”. In an EDHMM or latent smooth model, the former can be represented as a regular (semi-Markov) stage, while the latter can be treated as fixed duration micro-states. Let 𝒮_*F*_ ⊂ 𝒮 denote the set of flat states. Considering that the expected value of the signal at each electrode is assumed to be zero and that signals from different electrodes are assumed to be conditionally independent, it is reasonable to assume that

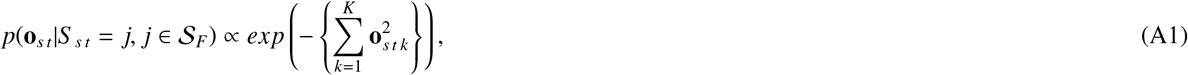

i.e., that the density of observing **o**_*s t*_ during a flat state is proportional to the negative sum of squares (taken over the *K* electrodes) of the EEG amplitudes (Anderson et al., 2016).

Matters are slightly more complicated for bump “states”: in contrast to flat states, for which the expected value is simply zero for all electrodes and at all times, the expected value now depends on the residual duration *τ*_*t*_ = *d*^′^ of bump state *j* and can be different for different electrodes. Specifically, considering that bump states are assumed to have fixed duration 𝒟^′^, the expected value at electrode *k* is equal to *s*(*d*^′^)*µ*_*j k*_ when the latent process is in state *j* S_*F*_ with residual duration *τ*_*t*_ = *d*^′^ and *d*^′^ ∈ {1, …*D*^′^}. *s*(*d*^′^) returns the amplitude of a half-sine of duration 𝒟^′^ time-locked to *d*^′^ = 1 at time-point *D*^′^ − *d*^′^ + 1 and *µ*_*j k*_ corresponds to the aforementioned weight of the half-sine added to the signal of electrode *k* when the latent process enters state *j* (Anderson et al., 2016). Note, that the value returned by *s*(*d*^′^) depends on the sampling frequency at which data was collected and the assumed frequency of the sine: Anderson et al. (2016) assumed a 10 Hz frequency for the sine, resulting in a 50 ms duration of bump states. Assuming that data is collected at a sampling frequency of 100 Hz, 𝒟^′^ = 5 and *s*(5) = 0.309, *s*(4) = 0.809, *s*(3) = 1.00, *s*(2) = 0.809, *s*(1) = 0.309 (Anderson et al., 2016).

Based on these considerations, the density of observing **o**_*t*_ conditional on the residual duration of state *j* ∉ S_*F*_ can then be defined as *P*(**o**_*s t*_|*S* _*s t*_ = *j, τ*_*s t*_ = *d*^′^, *j* ∉ S_*F*_) ∝

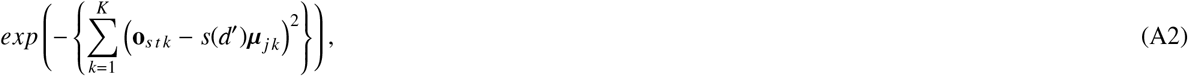

for all *d*^′^ 1, …*D*^′^. Fortunately, access to this density is sufficient to compute the forward variables defined in Equations (6) and (7). Note, that this will still require specification of the missing normalization constant to convert the sum-of-square terms on the right-hand side of Equations (A1) and (A2) into actual densities. In practice, this can be achieved by assuming a known distribution for the squared deviation of the amplitude at a given electrode from the expected values (i.e., 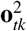 and **o**_*tk*_ − *s*(*d*^′^)*µ* ^2^). Plausible choices are Exponential, Chi-square, Gamma, or Normal distributions (see Anderson et al., 2016, for a more detailed discussion of plausible distributions). Finally, note that *µ*_*jk*_ can readily be parameterized by an additive mixed model as shown in Equation (3). Notationally, all that remains necessary to use the Emission model by Anderson et al. (2016) as part of a latent smooth is to replace *µ*_*j k*_ with *µ*_*s t j k*_ in Equations (A1) and (A2).

### ar-1 Model of the Deviations

Assuming a Gaussian model also enables simple auto-correlation (ar) models of the deviations (e.g., Wood, 2017), let 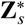 denote the true sequence of bump and flat states visited on trial *s*. Assuming that the deviations of the amplitude at a given electrode *k* from the expected values (i.e., **o**_*tk*_ − 0 and **o**_*tk*_ − *s*(*d*^′^)*µ*_*jk*_) are normally distributed with zero mean and variance *σ*^2^, the conditional density of all *T*_*s*_ amplitude recording **o**_*s k*_ collected on trial *s* at electrode *k* is multivariate normal, since 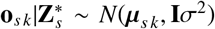. *µ*_*s k*_ is the vector of expected values containing zeros for every time-point 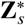 indicates to be in a flat state and the correct “bump” weights for electrode *k* and state *j* for every time-point 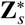 indicates to be in bump state *j*. As discussed in the main paper, here *µ*_sk_ is specified in terms of an additive (mixed) model. Thus, we write 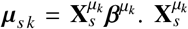 is obtained by stacking rows of zeros (for observations corresponding to flat states) and rows from the state-specific model matrices 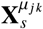 (for observations corresponding to bump state *j*). Similarly, 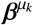 is obtained by concatenating all of the state-specific coefficient vectors 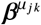 for all bump states. Since flat states are assumed to have an expectation of zero, no coefficient vectors are associated with them.

To include an ar model of the deviations we now substitute the covariance matrix **V** of an ar process (see Wood, 2017, for details on how to form **V**) for **I** and instead assume that 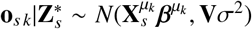. Applying standard results about linear transformations of normal random variables, we get 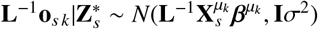, where **L**^−1^ is the inverse of the Cholesky factor **L** of **V** = **LL**^⊤^ (e.g., Wood, 2017). If 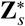 were known, estimating 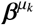 for an ar model could thus proceed in the same way as before – requiring only that the transformations of **o**_*s k*_ and 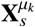 are computed once, before parameter fitting. Typically, 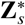 will however not be known so that the transformation cannot be computed in advance. Importantly, for lower-order ar models (e.g., of lag 1, resulting in an ar-1 model) computing **L**^−1^**o**_*s k*_ and 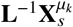 corresponds to a simple an efficient re-weighting of rows in **o**_*s k*_ and 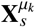 (Wood, 2017). As such, for lower-order ar models it remains possible to compute this re-weighting dynamically, while evaluating the forward variables, without too much computational overhead. Attention has to be paid to computing the expected value during the onset of a bump state: assuming an ar-1 model and a strict left-to-right HsMM with supra-diagonal transition matrix, as suggested by Anderson et al. (2016), the previous expectation, for the last observation emitted during a flat state, will always be zero. Additionally, when computing the gradient of the log-likelihood, it will be necessary to carefully consider cross-state dependencies of the emission densities: again assuming an ar-1 model, the density of the first observation of the current (flat) state will depend on the coefficients of the preceding (bump) state.

## Appendix B

### Partial Derivatives of Densities and Probabilities Required for Gradient Evaluation

In this appendix we show how the partial derivatives of the different densities/probabilities, all parameterized through different linear predictors, involved in Equations (12), (13), and (14) of the main paper can be obtained via transformation. This relies on the general strategy described by Wood et al. (2016). Let *β*_*r*_ again denote a coefficient included in the total set of coefficients *β* required by the models of all parameters. Then, assuming that *β*_*r*_ is involved in the model of parameter *µ*_*j*_,

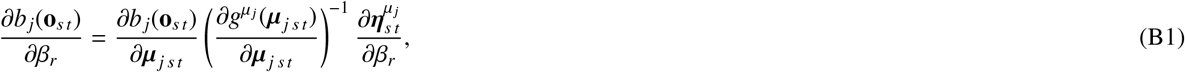

where 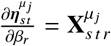 with 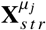 indexing column r in the row of model matrix 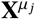 associated with time-series *s* and time-point *t*. Fortunately, the derivatives of the densities with respect to their parameters (i.e.,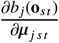) are available for many distributions – including the multinomial distribution, which enables computation of the derivatives involving *π* and Δ (e.g., Wood et al., 2016).

Computation of the derivatives of the state sojourn time densities 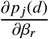 with respect to *β*_*r*_ might require additional work. If we assume a continuous distribution (i.e., ℱ^*d*^) for state durations *d*, then the derivative can be computed exactly as shown in Equation (B1). However, it is common to normalize the densities to ensure a proper probability mass function over the discrete interval {1, *D*}, in which case 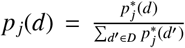, with 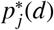 corresponding to the density of a continuous distribution (i.e., ℱ^*d*^; see also Yu, 2010). In that case, the required derivative has to be computed as shown in Equation (B2) – note that 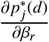 can again be computed as outlined above (i.e., Equation B1).

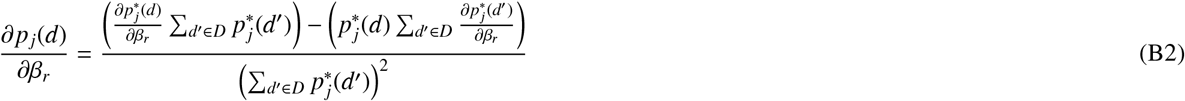

## Appendix C

### Subject-level Topography Estimates

**Figure C1.**
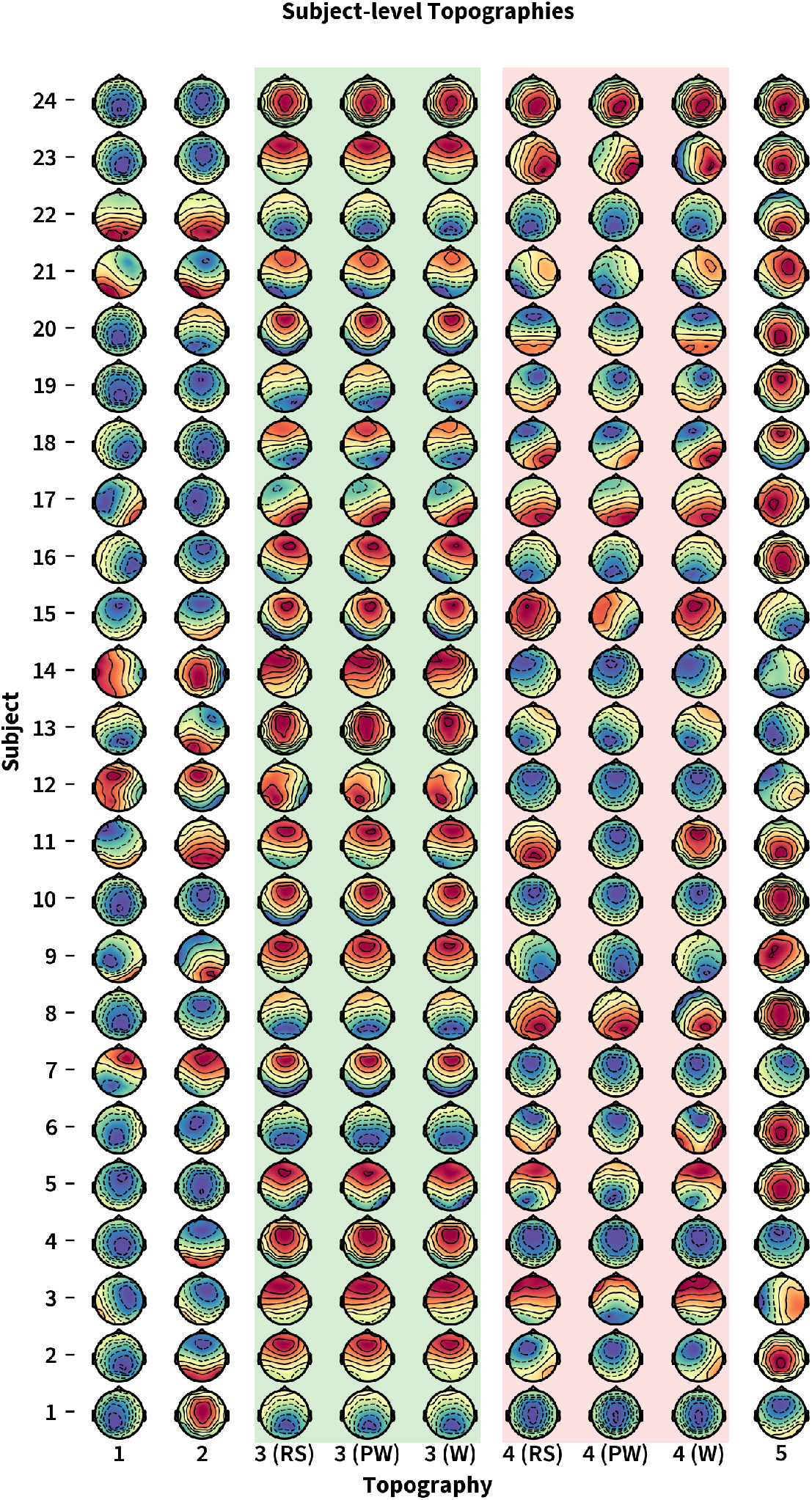
Subject-level Topography Estimates for all Word-types and Subjects. Figure C1 shows the subject-level estimates of the EEG topographies for all subjects and word-types. For the third and fourth topography (highlighted by color bars), separate group-level estimates were obtained per word-type (Words (“W”), Pseudo-words (“PW”), and Random Strings (“RS”)), which also results in word-type specific subject-level estimates for these topographies (although the same subject-specific difference is added to the group-level estimates).

## Appendix D

### Credible Intervals of the Difference in the Effect of Word-likeness

**Figure D1.**
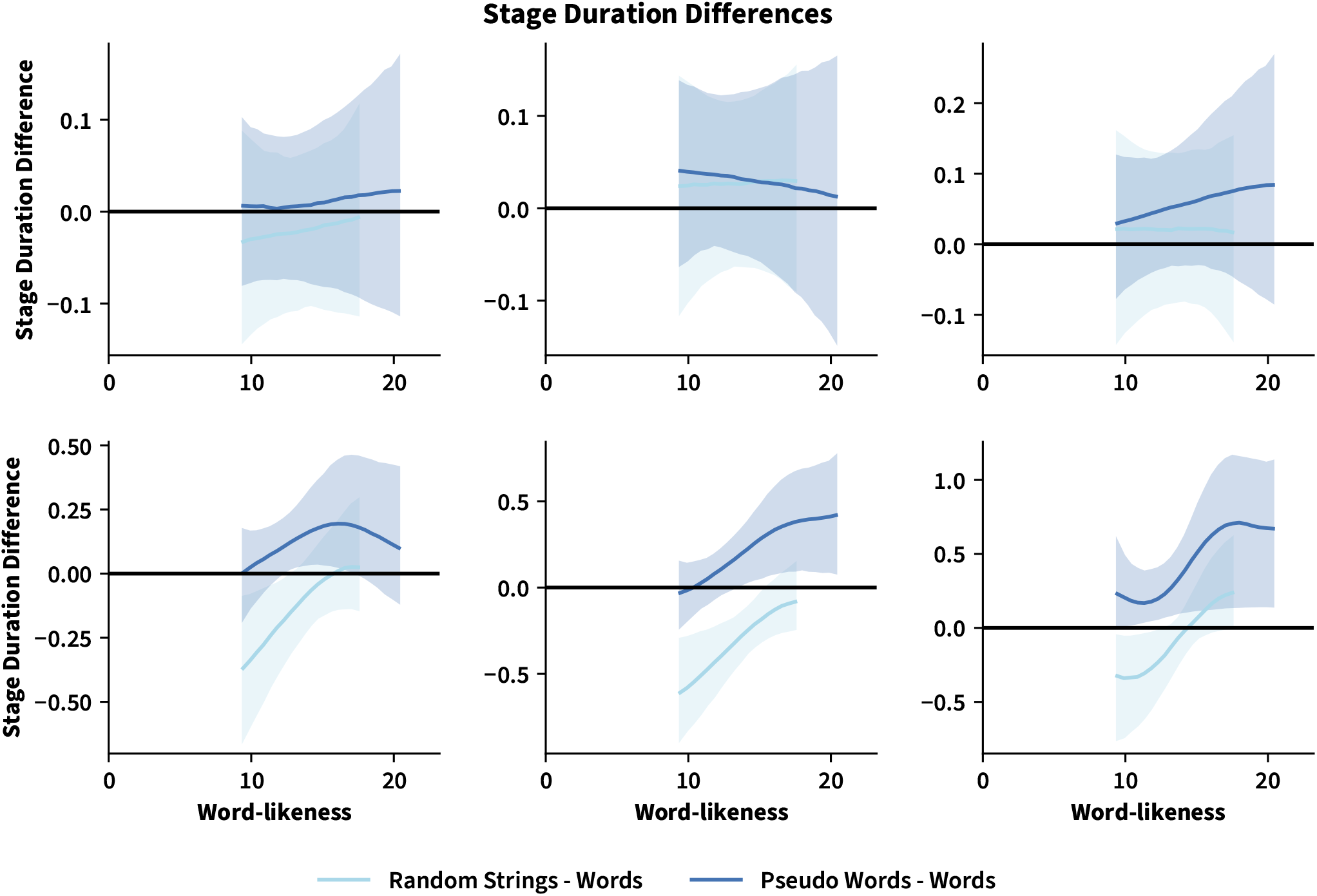
Credible Intervals of the Difference in the Effect of Word-likeness between non-words and words. Figure 7 shows an overview of the posterior mean estimate of the difference between the smooth functions *f*_*w*(*s*) *j*_(*wl*(*s*)), involved in the models of the mean parameter ν_*j*_ of the state sojourn times, between both types of non-words (random strings and pseudo-words) and words. Each panel corresponds to a different flat state, and visualizes the estimated difference smooth for both comparisons. Shaded areas are 95% posterior credible intervals for the difference. Evidently, later states show stronger differences in the effect of word-likeness between non-words and words. Importantly, the magnitude of the difference varies as a word-type specific function of word-likeness, word-type, and flat state – with the strength of the effect (measured by whether the credible interval includes zero or not) varying for different combinations of these three variables.

## Appendix E

### Residuals of Six Flat-State Multi-level Semi-Markov Model

**Figure E1.**
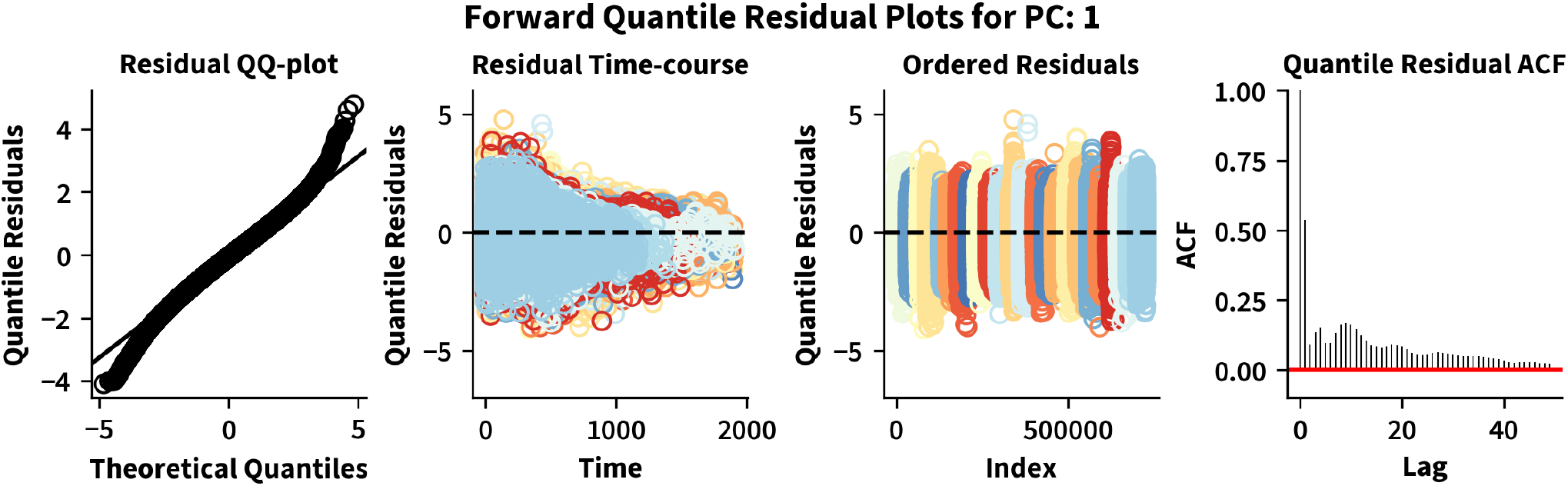
Residuals of Six Flat-State Multi-level Semi-Markov Smooth Model. Like Figures 3 and 4 in the main paper, Figure E1 shows different plots of the forward pseudo-residuals, but computed for the final multi-level semi-Markov Smooth model presented in the main paper. Notably, the forward pseudo-residuals of the multi-level model have only improved marginally. The main difference between the multi-level model and the group-level model (see Figure 4) is that the variance of the multi-level residuals differs less between subjects.

**Figure E2.**
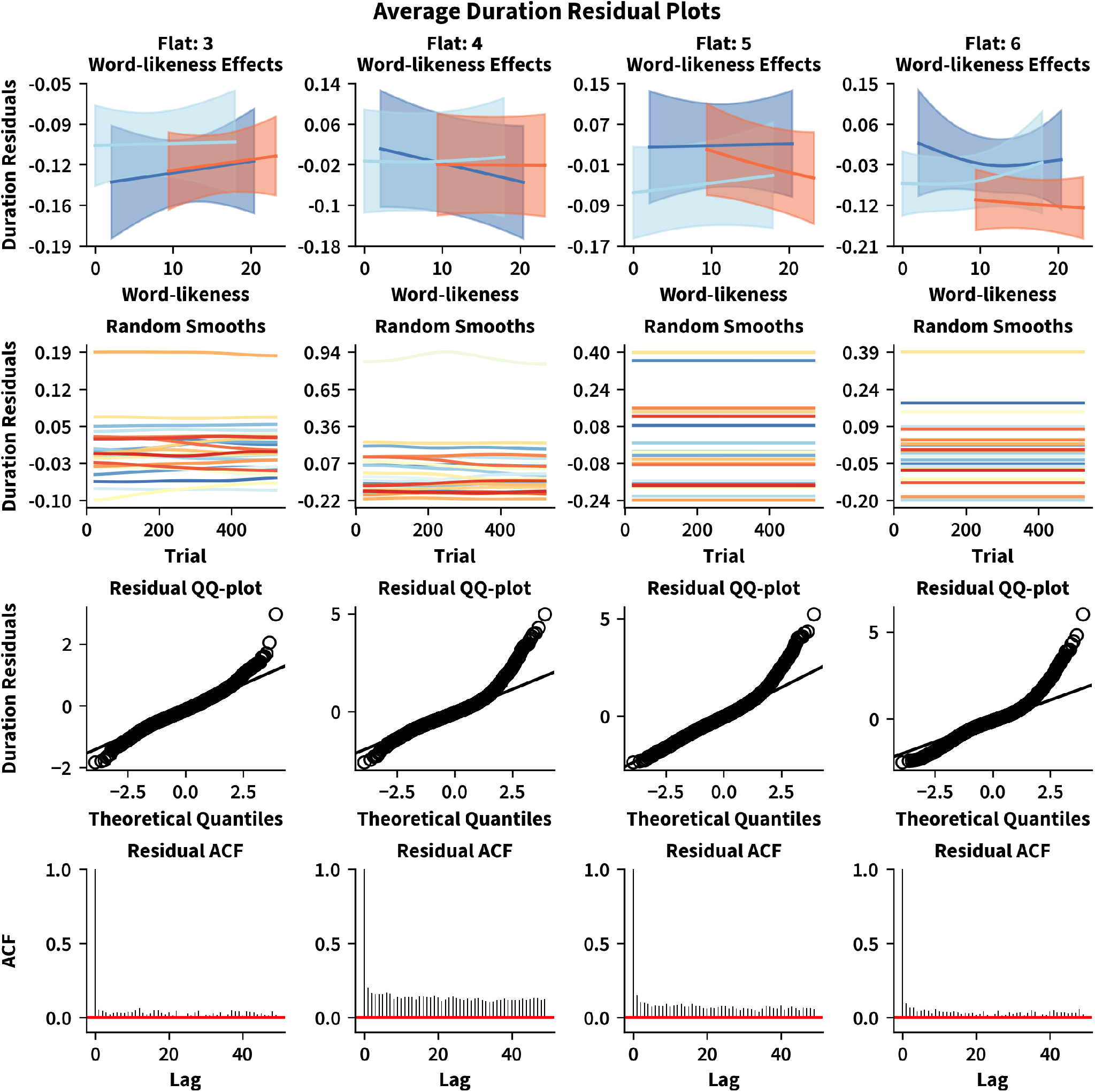
Average Flat-State Duration Residuals of Six Flat-State Multi-level Semi-Markov Smooth Model. Like Figure 5 in the main manuscript, Figure E2 shows plots of the average duration residuals for all flat states, but computed for the final multi-level semi-Markov Smooth model presented in the main paper. Note, that the average duration residuals now provide only week evidence of group-level effects of word-likeness or subject-specific changes in the expected residual over trials (note the y-axis differences when comparing with Figure 5). This is expected, considering that the multi-level model explicitly accounted for these effects. QQ-plots still show deviations from normality, but the residuals are far less correlated across different lags.

## Footnotes

1 See https://github.com/JoKra1/mssm.

2 Notably, Michelot (2025) discussed how to *approximate* explicit duration models with HMMs. In contrast, the algorithms presented here are exact.

3 So that 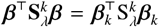, where 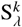 is any (semi-definite) precision matrix associated with the (improper) prior placed on *β*_*k*_ ⊂ *β*.

4 So 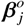 denotes the set of coefficients required by the models of all parameters (e.g., mean, scale, etc.) required to parameterize ℱ^*o*^

5 As an exception, this will not be necessary if Δ and *π* are set up in a way that guarantees that the Markov chain never re-enters the first state. Examples are strict left-to-right models for which *π*_1_ = 1 and Δ is a supra-diagonal matrix.

6 >The pre-processed data is openly available at https://github.com/JoKra1/HSMM_LD_EEG. Code to replicate the analysis and all figures is included in the Supplementary GitHub repository for this paper, available at https://github.com/JoKra1/SMSM_supplementary.

7 Corresponding to an alpha parameter of *α* = 2 if the Gamma distribution were to be defined in terms of an alpha and scale parameter.

8 There are always *M* −1 bump states and *M* flat states in the EDHMM defined by Anderson et al. (2016), since a topography is assumed to precede the onset of each process but the very first.

9 Considering the vast space of possible SMSMs for any given data-set, this will be the case for any automated selection strategy anyway. Careful consideration of existing theories and available experimental evidence to narrow down the potential number of states and effect structure will thus generally be advisable and typically increase the chances of a subsequent selection procedure to generate useful estimates.

